# Observation-Related Activity in Human Motor Cortex Increases with Effector Anthropomorphicity

**DOI:** 10.64898/2026.04.24.720491

**Authors:** Jacob T. Gusman, Zoe C. Beckman, Tyler S. Singer-Clark, Angelique C. Paulk, Anastasia Kapitonava, Tommy Hosman, Shane Allcroft, Alexander J. Acosta, Claire Nicolas, Daniel B. Rubin, John P. Donoghue, Carlos E. Vargas-Irwin, Leigh R. Hochberg

**Affiliations:** School of Engineering, Brown University, Providence, RI, USA; Carney Institute for Brain Science, Brown University, Providence, RI, USA; Department of Neuroscience, Brown University, Providence, RI, USA; VA Center for Neurorestoration and Neurotechnology, VA Providence Healthcare, Providence, RI, USA; Center for Neurotechnology and Neurorecovery, Department of Neurology, Massachusetts General Hospital, Boston, MA, USA; Department of Neurology, Harvard Medical School, Boston, MA, USA

**Keywords:** Human Motor Cortex, Mirror Neurons, Robotics, Anthropomorphism, Brain-Computer Interfaces

## Abstract

Neurons in motor cortex can be engaged not only in motor execution but also during observation of movements performed by other anthropomorphic agents (i.e. humans or monkeys). However, it is unknown how motor cortical neurons respond during observation of the range of assistive or prosthetic devices controlled by people using intracortical brain-computer interfaces (iBCIs). We recorded single-unit activity in the precentral gyrus while iBCI users viewed grasp-like movements performed by a spectrum of virtual effectors that included human, robotic, and hand-like dot stimuli. We found a relationship between neural modulation and effector *anthropomorphicity* (i.e. human-likeness) that existed on an ensemble-wide and individual neuron level, suggesting that human motor cortex activity incrementally increases in response to the visually observed agent’s human-likeness. Both solicited and spontaneous feedback from the participant indicated a relationship between neural activity and subjective assessments of anthropomorphicity, revealing a powerful contribution of context on observation-induced activity in motor cortex. The activity of motor cortex remained similar during attempted hand movements while different effectors were being observed, suggesting that intuitive external device control via iBCIs may not be overtly affected by the anthropomorphicity of the effector.

**SIGNIFICANCE STATEMENT:** The tendency for neurons in motor cortex to respond during movement observation has been proposed to underlie cognitive processes from motor learning and language development to empathy and theory of mind. Understanding how the motor cortex is engaged during observation of abstract and anthropomorphic agents informs our understanding of these processes and may guide development of neural prostheses which harness the activity of motor cortical neurons to restore lost neurologic function. Here we provide unique neuron-level evidence that human motor cortex activity is gradually modulated by how human-like an observed agent appears and moves. This finding advances our interpretation of “mirror” activity in the brain and could help guide the design of brain-controlled prostheses used by people with tetraplegia.

## INTRODUCTION

The ability to rapidly perceive, model, and predict actions performed by other humans is a fundamental capability of the human brain. No single finding in recent decades has shaped the conversation on this process as profoundly as the concept of “mirror neurons”. After their discovery in the monkey ventral premotor cortex^1^, mirror neurons were initially defined as any neuron that responds during both the execution and observation of biological movement^2^. Whereas intracortical recordings have confirmed the presence of similar responses among neurons in dorsal premotor^3,4^ and primary motor regions^3,5,6^ of the nonhuman primate, our understanding of mirror-like activity in human motor cortex has been largely limited to functional imaging studies^7–9^. However, with the development of intracortical brain-computer interfaces (iBCIs), chronic high-resolution recordings of human motor cortex are increasingly available, yielding an opportunity to not only study the effects of mirror-like modulation on motor decoding but also revisit fundamental questions about mirror neurons with the precision afforded by single-unit recordings.

A defining characteristic of mirror neurons is their biological “tuning” to the observation of human and monkey movements. But how does the brain determine what is “human”, and how does the motor cortex respond to the observation of artificial, humanoid effectors such as the prosthetic and assistive robot arms utilized by iBCI users? Human neuroimaging studies have evaluated mirror activity during robot observation by comparing observation of human movements to one or two robotic exemplars, resulting in contradicting assessments of whether or not the mirror phenomenon extended to robots^10–16^. The fact that each of these studies utilized different robotic exemplars that differed along multiple dimensions (color, shape, movement, etc.) offers the possibility that visual responses in motor cortex could depend on a continuous measure of “human-likeness” – an effect that would necessitate higher resolution recordings to ascertain.

Here, we studied the neural correlates of *anthropomorphicity* – operationally defined as the relative visual similarity of an effector to a typical human hand – in the left precentral gyrus of two iBCI clinical trial participants with tetraplegia. Participants viewed a spectrum of human, robotic, geometric, and dot animations, revealing an anthropomorphic *gradient* in population and individual neuron responses in the dorsal premotor cortex. We also found that visually-induced modulation can be affected by “top-down” variables: in a session with one participant, we demonstrate that his motor cortex only became engaged after he *realized* that a dot stimulus was meant to represent a hand. Finally, we evaluated neural activity patterns when participants observed virtual effectors while simultaneously performing attempted hand grips, indicating that the influence of anthropomorphicity in motor cortex may be significantly diminished in the presence of the motor signals that are used to operate iBCIs. Together, these findings provide the most detailed account to date of observation responses in motor cortex, contributing important context for interpreting prior neuroimaging studies on mirror neurons and offering insight into how abstract, high-level variables – such as anthropomorphicity – can be represented at the level of individual neurons.

## RESULTS

Two participants with tetraplegia, T11 (spinal cord injury, C4 AIS-B) and T17 (advanced ALS, ALSFRS-R = 0), had microelectrode arrays implanted in their left precentral gyrus (PCG) as part of their enrollment in the BrainGate2 clinical trial (**SI Appendix**). Participant T11 had two 96-channel microelectrode arrays (Blackrock Neurotech) surgically implanted in the “hand knob” area of the left PCG, which was identified by pre-operative magnetic resonance imaging (MRI). Participant T17 had six 64-channel microelectrode arrays implanted in motor- and speech-related areas along the left precentral gyrus. MRI/CT coregistration determined that locations of the two arrays in T11’s PCG correspond to Brodmann Area (BA) 6d and the six arrays in T17’s PCG correspond to areas 6d, 55b, and 6v (two arrays per area, **Figure 1A**; see **SI Appendix**). For this study, neural signals were recorded from arrays in areas 6d (T11 and T17) and 6v (T17), and custom software was used to identify single-unit potentials (see **SI Appendix**; **Tables S1-S2** list the number of trials and number of single units analyzed in each session).

**Figure 1.**
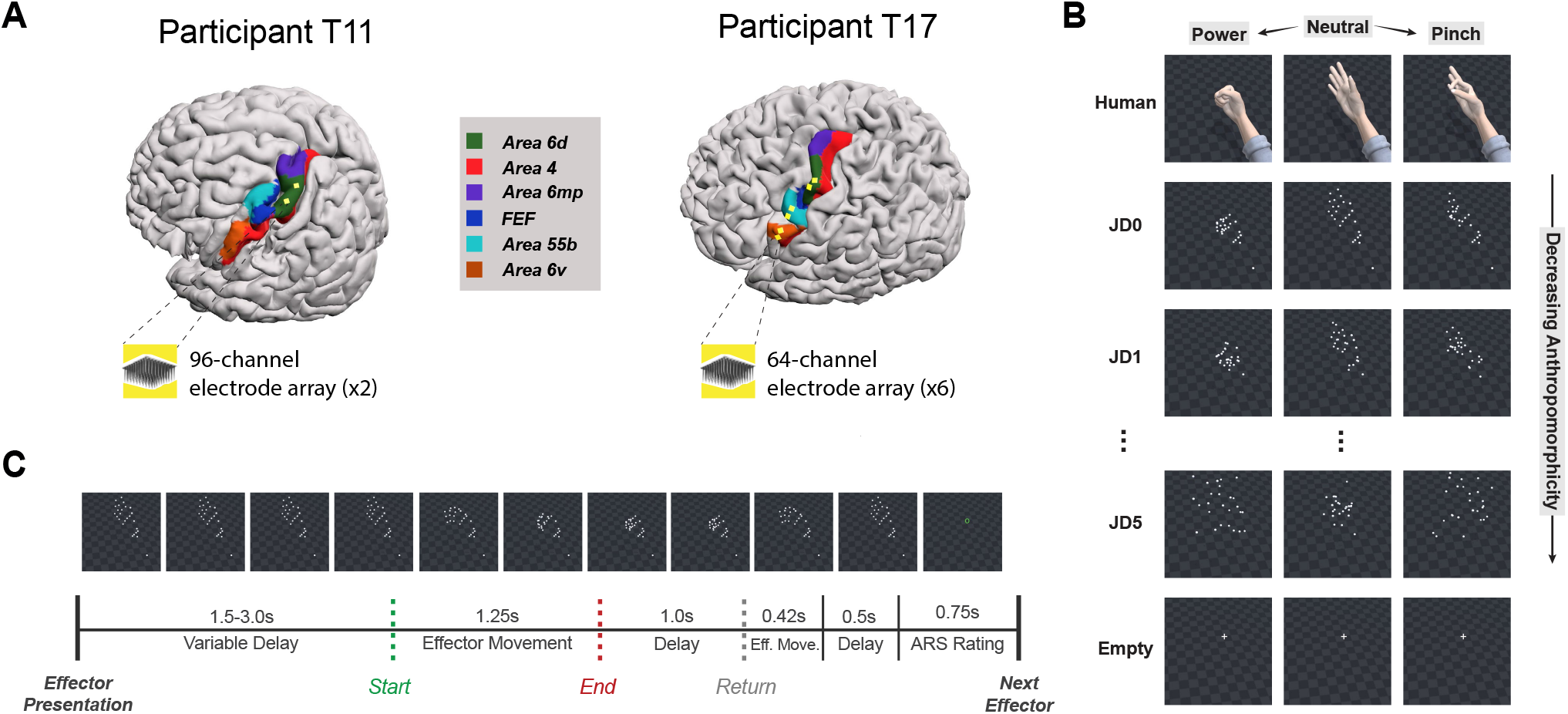
The Jittered Dots Task. (A) MRI-derived 3D brain reconstructions for participant T11 and T17 showing the implant locations of electrode arrays (yellow squares) in the left precentral gyrus. Colored shadings were created using cortical labeling from the Human Connectome Project (HCP) Atlas parcellation labels transformed into FreeSurfer space and then reconstructed into Blender (see **SI Appendix**). (B) Still frames of example stimuli from the Jittered Dots (JD) Task at the start (“Neutral”) and end of effector movements (“Pinch” or “Power”). In total, eight stimulus conditions were shown to the participant with half the trials showing a “power grip” movement and half showing a “pinch grip” movement. Additional still frames, videos, and written descriptions of all conditions can be found in **Figure S1, Video S1**, and **Table S3**, respectively. (C) Temporal sequence of each trial with still frames from an example JD0 power grip trial. A green circle was used at the end of each trial to indicate when the participant was to provide his anthropomorphicity rating using the anthropomorphicity rating scale (ARS, see **SI Appendix**).

### Ensemble Activity in Motor Cortex is Tuned to Effector Anthropomorphicity

Our first objective was to characterize the relationship between neural activity in motor cortex and anthropomorphicity. Specifically, does the motor cortex respond in a *binary* fashion – only responding to observed movement once a certain anthropomorphicity threshold is met – or is it gradual – displaying neural activity patterns that are ordered along an anthropomorphic spectrum? Human-likeness is a complex and often subjective metric influenced by a wide range of factors including physical appearance, context, and prior experience^17–19^. Therefore, we designed the Jittered Dots (JD) task, a novel dot stimulus paradigm that allowed us to systematically “jitter” the features of a hand-shaped dot animation to evaluate anthropomorphic tuning in a controlled, gradual manner (**SI Appendix**).

In the JD Task, participants were asked to passively observe 8 stimuli: a virtual Human Hand that was animated (Unity Technologies, San Francisco, CA) to perform “power” or “pinch” grip motions, a static cross (referred to as “Empty” herein), as well as 6 dot stimuli that were altered to span a spectrum ranging from obviously hand-like (JD0) to completely random three-dimensional motion (JD5) (**Figures 1B** and **S1**; see **Table S3** for stimuli descriptions; **Video S1**). Note that the JD Task stimuli were generated by gradually introducing changes to an original 3D human hand animation, resulting in a set of stimuli that are ordered by anthropomorphicity but do not necessarily represent a linear relationship in this respect. Except for the Empty condition and JD5, each stimulus had distinct “power grip” and “pinch grip” variations. We presented the stimuli pseudorandomly using the trial structure shown in **Figure 1C**. In addition to watching the screen, we instructed the participants to assess the hand-likeness of each stimulus and provide a numerical rating (1 through 4) at the end of each trial according to the Anthropomorphicity Rating Scale (ARS; **SI Appendix**). Participant T11 performed the JD Task across 3 session days and T17 performed one session of the JD task.

#### Ensemble firing rate gradually increases with stimulus anthropomorphicity

As a general measure of activation in each brain region, we calculated the percent increase in firing rate (FR) of each neural ensemble (6d_T11_, 6d_T17_, and 6v_T17_) during observation of each JD Task stimulus compared to baseline (**SI Appendix**). Increases in ensemble FR followed a pattern that was consistent with a *monotonic* relationship between stimulus anthropomorphicity and motor cortex modulation (**Figure 2A-B, Figure S2A**). Although most apparent in responses from 6d_T11_ (**Figure 2A**), directional (one-tailed) Wilcoxon Rank Sum (RS) tests revealed that there were “intermediate” responses between the most anthropomorphic (e.g. Human, JD0) and least anthropomorphic (e.g. JD5, empty) in responses from all ensembles (see **Table S4**). For example, the ensemble FR increase when T11 viewed JD1 was significantly less than that of JD0 (*p* = 8.74 × 10^-6^) and significantly greater than JD2 (*p* = 2.40 × 10^-25^). Responses in 6d_T17_ (**Figure 2B**) during JD0 observation were less than Human observation (*p* = 2.22 × 10^-5^) and greater than JD5 observation (*p* = 0.028). Responses in 6v_T17_ (**Figure S2A**) during JD2 observation were less than JD1 observation (*p* = 0.024) and greater than JD4 observation (*p* = 0.031). Furthermore, the trend was consistent across the spectrum, with no instances of a less anthropomorphic stimulus producing a significantly greater response than a more anthropomorphic stimulus (RS tests, *p* > 0.05; **Table S4**).

**Figure 2.**
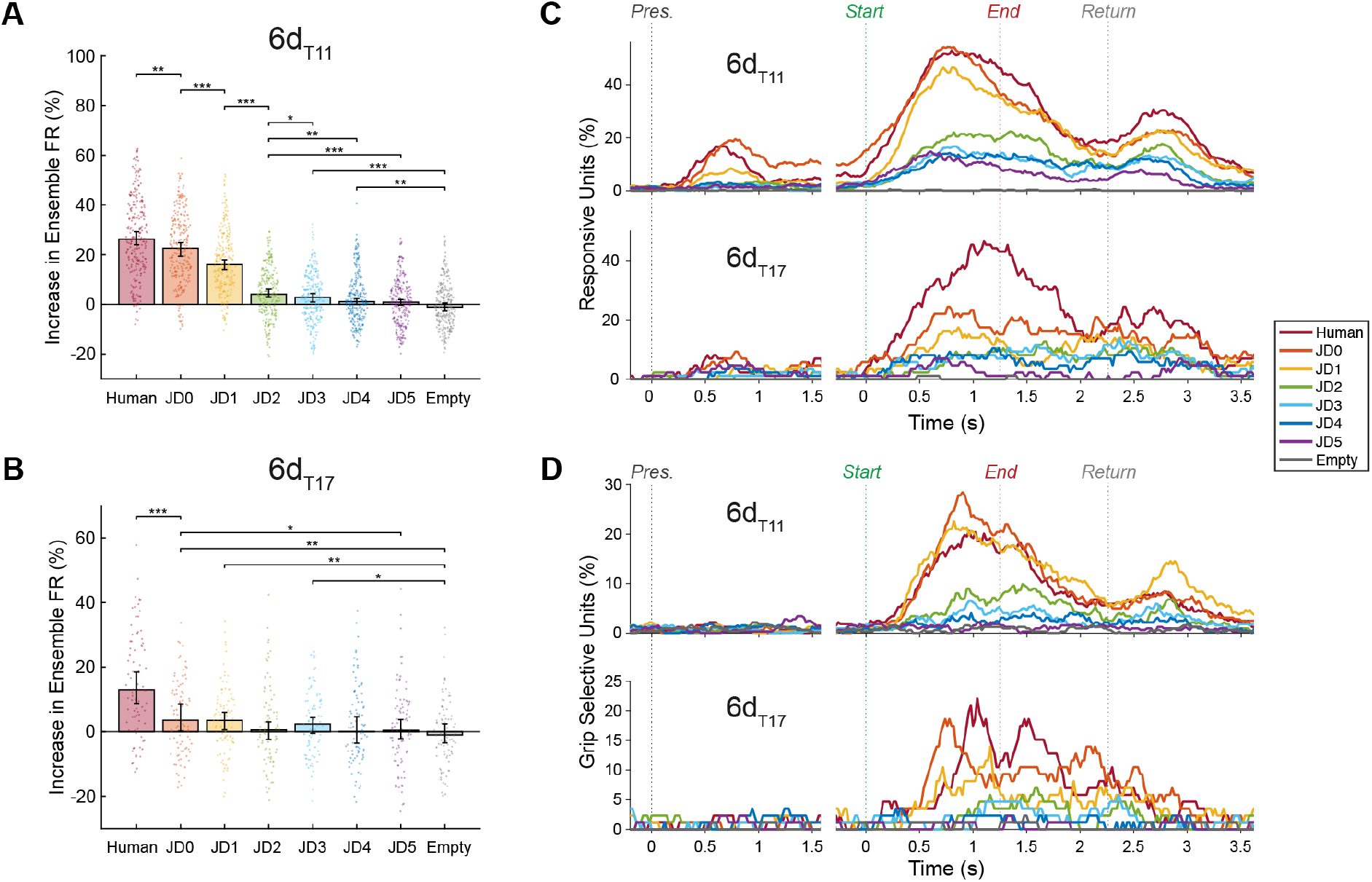
Modulation of the Motor Cortical Ensemble During the Jittered Dots Task. (A-B) Increases in ensemble firing rate (FR) in area 6d (see **Figure S2** for T17-6v results) during passive observation of each of the eight stimuli presented during the JD Task. Each point in the swarm plots represents the increase of the mean firing rate of the recorded ensemble (calculated on a 1.5s window beginning 200ms after Start) as a percentage of the block baseline FR for one trial. Block baselines were calculated as the mean FR of all the Empty trials in a given block. Bar heights denote median increase in ensemble FR; error bars denote 95% confidence intervals (CIs) of the median. Outlier trials outside of 1st-99th percentile range were omitted from each swarm plot but included in analyses. Horizontal bars indicate significant decreases between conditions (Wilcoxon Rank Sum, one-tailed). Asterisks indicate significant *p*-values (*, *p* < 0.05; **, *p* < 0.01; ***, *p* < 0.001). Nonsignificant (*p* > 0.05) and redundant comparisons are omitted (e.g. Human > JD1 is not shown because Human > JD0 and JD0 > JD1). *P*-values from all pairwise comparisons appear in **Table S4**. (C) Percent of units responsive to the task over the course of the trial period. A unit was considered responsive to a stimulus in a given window if a significant difference (Kruskal-Wallis test, p < 0.01) in firing rate was found when comparing its responses across the power grip condition, the pinch grip condition, and the baseline (Empty/Relax) condition (see **SI Appendix**). (D) Percent of units selective for grip type. Units were considered grip selective in a given time window if they had significantly different (RS test, *p* < 0.01) FRs between power and pinch conditions. Note different axis scales in A vs B, C(top) vs C(bottom), and D(top) vs D(bottom) panels, resulting from relatively smaller increases in FR (B) and fewer responsive/selective units (C, D) in T17’s recorded neural activity compared to T11’s.

#### Proportion of neurons activated during effector movement observation depends on anthropomorphicity

For a more detailed understanding of the timing of neural activation during the JD Task, we evaluated the proportions of *responsive* and *grip selective* units (presumptive neurons) over the course of the trial period. We categorized a unit as *responsive* to a given JD condition (or *effector*) if there was a significant difference in FR (Kruskal Wallis (KW) test, *p* < 0.01) among its responses during the power trials of that effector, the pinch trials of that effector, and the baseline condition (“Empty (relax)”). We categorized a unit as *grip selective* during observation of a given effector condition if it had a significantly different FR between power and pinch trials (RS test, *p* < 0.01). We then calculated the percentage of units that were responsive and percentage of units that were grip selective in a 300ms sliding window shifted in 20ms increments across the trial epoch (see **SI Appendix**).

As expected, the largest proportion of responsive units and grip selective units was observed during the Effector Movement (**Figure 1C**) period of the trial, with peaks occurring around one second after the start of stimulus movement (**Figure 2C-D, Figure S2B-C**). For both participants, the maximum number of responsive units in 6d (6d_T11_, 6d_T17_) during this period was related to anthropomorphicity of the stimulus, with two minor exceptions: in 6d_T11_ JD0 elicited a slightly higher peak in responsive units (54.2%) compared to Human (52.9%) and JD5 (14.9%) was slightly higher than JD4 (14.2%). Interestingly, we observed a second, smaller, anthropomorphically arranged ramp-up in unit responsiveness after the end of the tested movement period as well. This could have resulted from observation of the “Return” movement of the effector that occurred at time 2.25s to 2.68s. In 6v_T17_, although there were similar peaks in responsiveness during both the Effector Movement and Return phases of the trial, the proportions of responsive units in 6v_T17_ did not appear to correspond with an anthropomorphic arrangement (**Figure S2B**).

Unit grip selectivity in area 6d also appeared to correspond with stimulus anthropomorphicity (**Figure 2D**). In 6d_T11_, although JD0 (28.5%) and JD1 (22.6%) yielded a greater proportion of grip selective units than Human (20.7%), the results of JD0-JD5 were consistent with their anthropomorphic ordering (JD2: 9.9%, JD3: 6.5%, JD4: 4.0%, JD5: 2.2%). Similar to unit responsiveness in 6v_T17_, grip selectivity results in 6v_T17_ were not obviously anthropomorphically arranged (**Figure S2C**), with the highest percentage of grip selective units exhibited during JD1 (20.0%) and JD2 (16.4%) observation, followed by Human (10.9%), JD0 (10.9%), JD4 (10.9%), JD3 (7.3%), JD5 (3.6%), and Empty (3.6%).

#### Gradual changes to ensemble spiking patterns reveal anthropomorphic gradient in responses to dot stimuli

While spike rates alone encode some information, additional insight is gained by taking into account the precise timing of individual spikes^20^. We hypothesized that spike train patterns may also reveal information related to the anthropomorphicity of observed visual stimuli. Using spike train similarity space (SSIMS) analysis (see **SI Appendix**), the firing pattern of the entire ensemble of simultaneously recorded neurons is projected as a single point in a latent space representation, and relative similarities of neural activity in each trial are described by their relative distances in the SSIMS projection. Although the data exist in a high dimensional space, it is possible to visualize the relationships between activity patterns using two-dimensional plots (e.g. **Figures 3A**).

**Figure 3.**
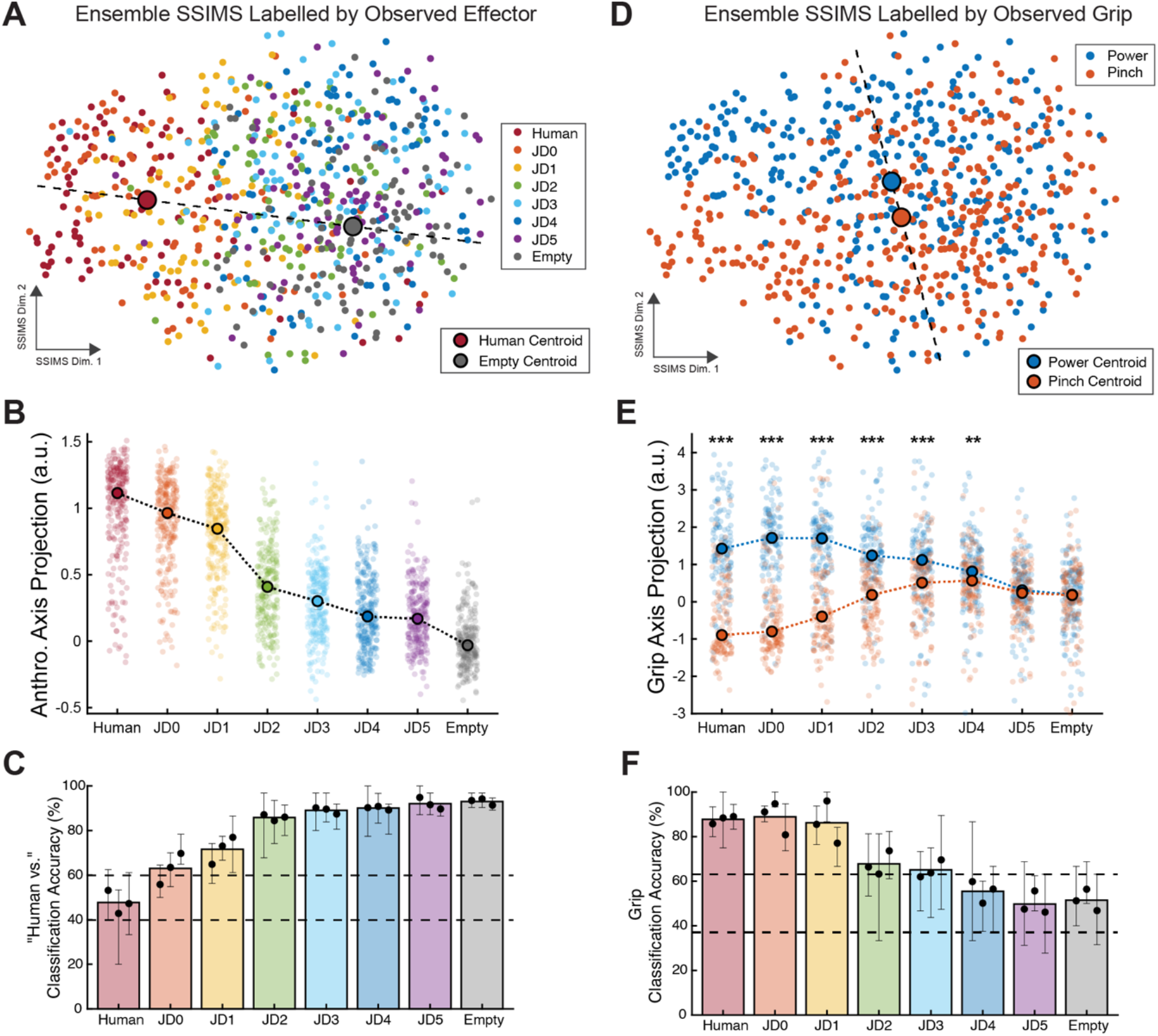
Ensemble Firing Patterns Reveal an Anthropomorphic *Gradient* in Spike Train Similarity Latent Space. (A) 2D spike train similarity space (SSIMS) plot of all trials from an example JD Task session (T11, trial day 412) colored by stimulus condition. Each marker represents the firing patterns of the entire ensemble of recorded neurons during a single trial, and the distances between markers represent the relative similarity between spike train patterns (**SI Appendix**). The large red circle indicates the centroid of the Human Hand trial data, and the large gray circle indicates the centroid of the Empty trial data. An anthropomorphic ordering can be observed along the “axis” connecting the centroids of the Human and Empty data. (B) Projections of the 10D SSIMS representations onto the “anthropomorphicity axis” for each stimulus condition. Anthropomorphicity axes for each session were defined by the line connecting the SSIMS centroids of the Human and Empty conditions. Trials from all three of T11’s JD Task sessions are shown. Connected dots represent medians for each condition. (C) Classification accuracies when a decoder is trained to distinguish between the Human Hand data and each of the other stimulus conditions. Black dots indicate mean classification performance after 5-fold cross validation for each session. Bar heights denote the mean over all sessions. Error bars denote the range of classification accuracies derived from the cross-validation process for each session. Dashed lines represent the 95% confidence interval of the chance distribution (see **SI Appendix**). Higher classification accuracies can be interpreted as indicative of decreased overlap between condition clusters in the 10D SSIMS representation, whereas accuracies that fall within the 95% confidence interval of the chance distribution describe conditions that have spike trains that cannot be easily distinguished from one another. (D) Same as (A), with trials color coded by grip type. Centroids for all power and pinch trials are shown with a line showing a 2D “grip axis” in the data. (E) Projections of the 10D SSIMS representations onto the “grip axis” for each stimulus condition. Grip axes for each session were defined by the line connecting the SSIMS centroids of all power and pinch conditions. Trials from all 3 of T11’s JD Task sessions are shown. Error bars represent the medians and 95% confidence intervals for each condition. Significant differences (RS test, two-tailed) between power and pinch trials are denoted by asterisks (*, *p* < 0.05; **, *p* < 0.01; ***, *p* < 0.001). (F) Same as (C), but showing classification performance of decoders trained to distinguish between power and pinch trials for each condition.

In 6d_T11_ and 6d_T17_, spike trains from anthropomorphically adjacent conditions were reliably similar to each other and appeared to self-organize according to anthropomorphicity in the 2D SSIMS projection (**Figure 3A, Figure S3A**). By projecting the 10D data to a line connecting the centroids of the Human and Empty conditions, we defined an “anthropomorphicity axis” (**SI Appendix**) that revealed that spiking patterns become less reflective of a human-like effector as dot stimuli become more random (**Figure 3B, Figure S3B**). To further quantify this trend, we created classifiers comparing the cluster of Human trials to the clusters of each of the other conditions (see **SI Appendix**) and found that classification accuracy (i.e., the ability to distinguish from “Human”) monotonically increased as the compared condition became less anthropomorphic (**Figure S3C, Figure S3C**). Similar to our analyses on 6v_T17_ firing rates (**Figure S2**), SSIMS analyses of 6v_T17_ spike trains did not demonstrate an obvious relationship with anthropomorphicity in area 6v (**Figure S4**).

Also evident in the SSIMS projection of 6d_T11_ spike trains was a dependence on anthropomorphicity with respect to grip type (**Figure 3D**). By projecting the data onto a “grip axis”, we saw that power and pinch spike trains become gradually more similar as the stimuli become less anthropomorphic (**Figure 3E**). Grip classification of 6d_T11_ SSIMS data confirm that distinguishing between neural activities of the two grip types was quite easy for Human, JD0, and JD1 (>85% accuracy), but gradually more difficult as stimuli become less anthropomorphic (**Figure 3F**). Although spike trains in 6d_T17_ and 6v_T17_ during power and pinch trials were separable when projected to a grip axis in SSIMS (**Figure S3E, Figure S4E**), there was not a clear relationship between anthropomorphicity and grip separability in the SSIMS projected T17 data (**Figure S3F, Figure S4F**).

### Individual Neurons in Motor Cortex are Tuned to Effector Anthropomorphicity

The decrease in ensemble FR associated with anthropomorphicity in area 6d (**Figure 1A-B**) could be caused by two phenomena at the single-unit level: the activation of fewer neurons in the ensemble or decreases in activity among already active neurons (or both). Our analysis of neuronal responsiveness (**Figure 1C**) suggests that fewer neurons become activated by less anthropomorphic stimuli. However, we also found that the responses of single units during the JD Task exhibited anthropomorphic “tuning” that could have also contributed to ensemble-wide modulation in the motor cortex (e.g. **Figure 4**, unit 68).

**Figure 4.**
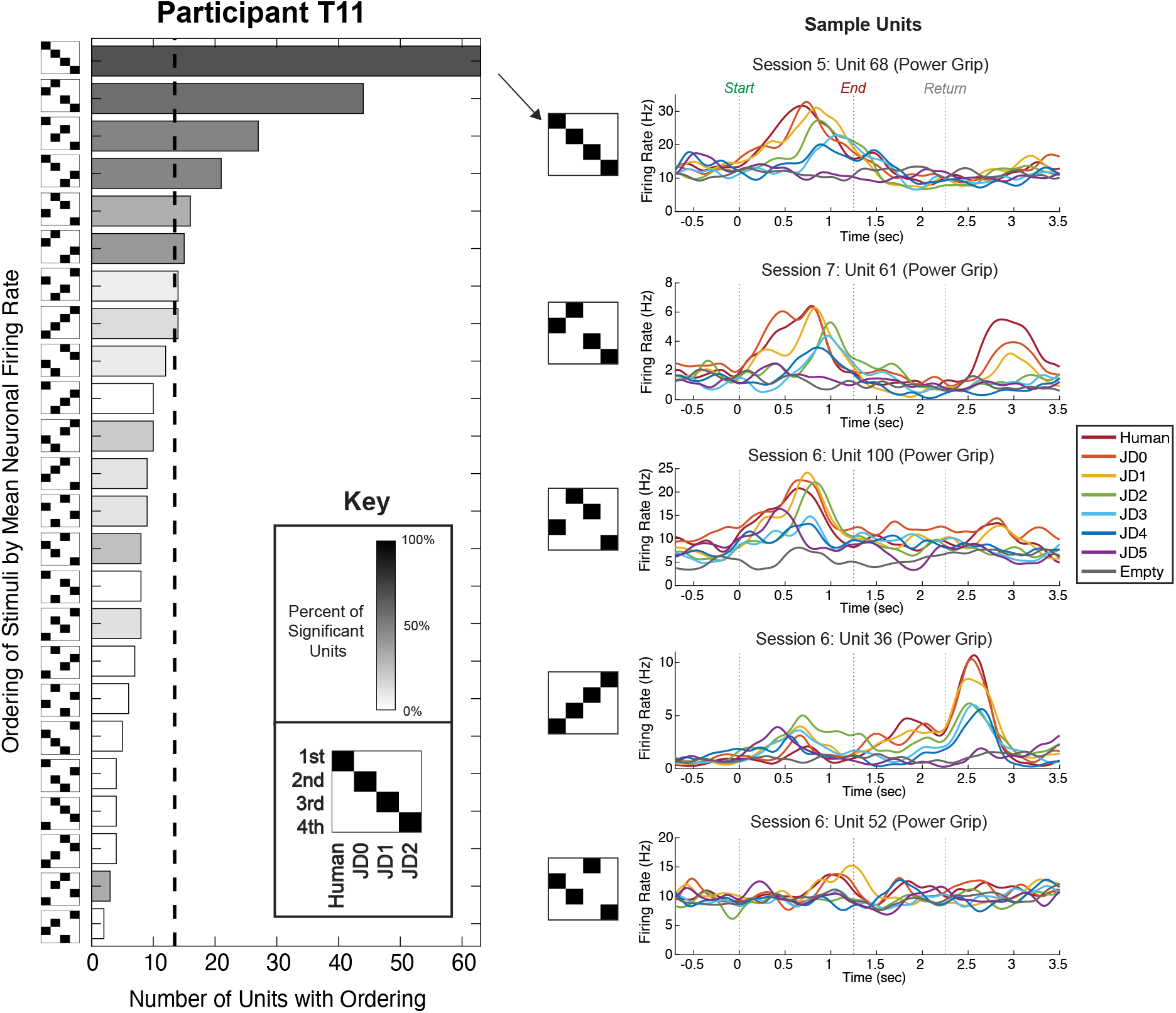
Single-Unit Firing Rates Reflect Anthropomorphic Tuning. Summary of the types of neurons recorded from participant T11 categorized by their relative responses to Human, JD0, JD1, and JD2 stimuli (“power” trials only; for T11 “pinch” trials see **Figure S5**; see **Figure S6** for T17 results). For each unit, responses to each stimulus condition were ranked according to the resultant mean firing rate during the 200-1700ms window following the start of stimulus movement. Bar plot lengths represent the number of neurons that had a particular ordering of mean firing rates. All 24 possible orderings are shown. The vertical gray dashed line reflects the number of neurons that would have a particular ordering by chance (∼13.5). The color (grayscale) of each bar represents the percent of units within a category that have significantly different firing rates between two or more conditions (Kruskal-Wallis, *p* < 0.01). Peristimulus time histograms for five sample units are shown.

For each neuron, we calculated its average FR across all trials for each stimulus condition and assigned the neuron a stimulus “ordering” in accordance with the magnitude of its response to each stimulus. For example, if a neuron on average fired at 10 Hz during Human trials, 8 Hz during JD0, and 5 Hz during JD1, its “ordering” with respect to those 3 conditions would be the following: Human (1st), JD0 (2nd), JD1 (3rd). For simplicity, we focused on the ordering of responses to “power” trials of the Human, JD0, JD1, and JD2 stimuli, which produced the most distinct differences in ensemble modulation among the effector conditions (similar results were obtained for pinch trials, see **Figure S5**). Across all ensembles, the most common ordering of responses to the stimuli was the anthropomorphic one, in which a neuron’s FR was highest for Human, followed by JD0, JD1, and JD2 (**Figure 4, Figure S5, Figure S6**). For example in 6d_T11_, of the 323 neurons assessed (combined across all three T11 JD Task sessions), 63 (20%) had anthropomorphically ordered mean FRs, which was nearly five times what would be expected by chance (13.5 units or 4.2%; **Figure 4**). Of these neurons, 43 were determined to have significantly different FRs between at least two of these conditions (KW test, *p* < 0.01), representing a far greater proportion (68.3%) than that of any other ordering. This suggests that not only were anthropomorphically ordered neurons more common, but they also produced more robust differences in FR between conditions.

The next most common ordering in 6d_T11_ (n = 44, 13.6%) was similar to the strictly anthropomorphic ordering but switched Human and JD0 positions. The relative ordering of JD0, JD1, and JD2 was unchanged, however. Indeed, 39% of all neurons (n = 127), and about two-thirds of all neurons with significantly different FRs (n = 73, 64%) had mean FRs that were anthropomorphically ordered for these first three jittered dot stimuli (see also **Figure S5**).

### Relationship Between Ensemble Activity and Anthropomorphicity Rating

By having participants provide their impressions of the anthropomorphicity for each stimulus via the Anthropomorphicity Rating Scale (ARS, see **SI Appendix**), we were able to relate their subjective assessments with objective gradations in the JD stimuli and their associated neural responses. As expected, participants consistently rated the Human stimulus a “4” (T11: 238 of 245 trials, T17: 89 of 89 trials) and rated the Empty stimulus a “1” (T11: 246 of 248 trials, T17: 90 of 90 trials). For both participants, the average ratings of the JD stimuli were ordered anthropomorphically from JD0 to JD5 (**Figure 5A**), confirming that our assumed anthropomorphic ordering based on the relative congruence of the jittered dot array with the original human hand model was consistent with the participants’ subjective assessments. However, there were some differences in how participants distributed their ARS ratings. For example, T11 rated Human (mean ARS: 3.9) and JD0 (3.8) very similarly, whereas T17 categorized Human (4.0) as seemingly a separate tier above JD0 (3.0) and JD1 (2.7).

**Figure 5.**
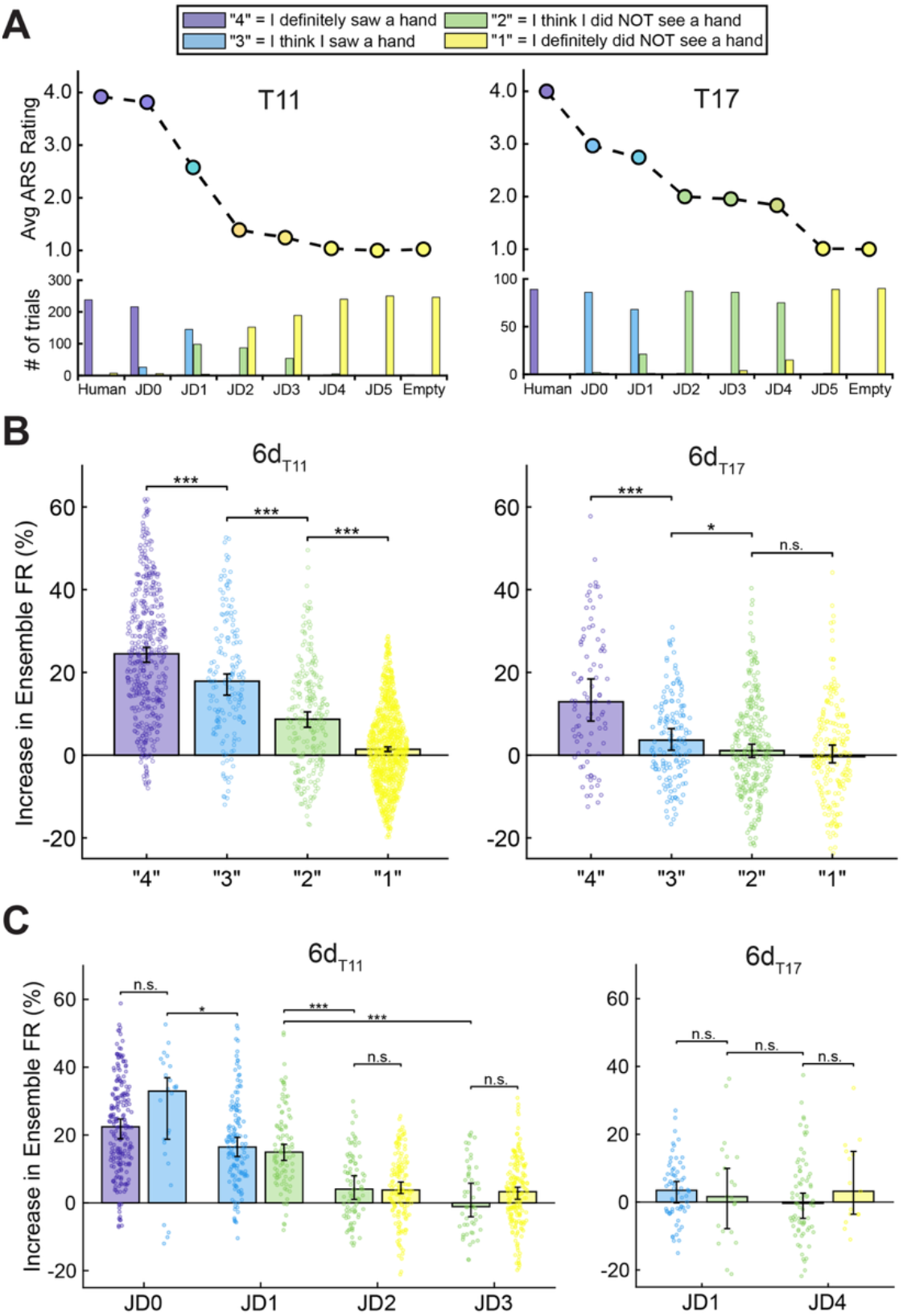
Effects of Participant ARS Rating on Ensemble Firing Rate. (A) Counts of ARS ratings given by each participant for each stimulus condition during the JD Task (bottom) and mean of ratings for each condition (top). (B) Increases in 6d ensemble firing rate (FR) for all trials, organized by the ARS rating given by each participant after each trial. Swarm plots, bar plots, and error bars are presented as in **Figures 2A-B**. Horizontal bars indicate significant decreases between conditions (Wilcoxon Rank Sum, one-tailed). Asterisks indicate significant *p*-values (*, *p* < 0.05; **, *p* < 0.01; ***, *p* < 0.001). Redundant comparisons are omitted (e.g. “4” > “2” is not shown because “4” > “3” and “3” > “2”). (C) Increases in 6d ensemble FR of trials for each stimulus condition, according to the given ARS rating. Only stimulus conditions that had more than 15 trials of more than one rating are shown. Swarm plots, bar plots, and error bars are presented as in **Figures 2A-B**. Horizontal bars indicate significant differences between trials of different stimulus conditions but the same assigned ARS ratings (Wilcoxon Rank Sum, two-tailed). No significant differences were found between trials of the same stimulus but different ARS rating.

Similar to the monotonic pattern of ensemble modulation that resulted from comparing the JD stimuli (**Figure 2A-B**), when trials were organized by their given ARS rating, we see a gradual decrease in ensemble FR corresponding to lower ARS ratings (**Figure 5B, Figure S7A**). However, because the objectively-defined anthropomorphicity gradient represented by the JD stimuli is correlated with the subjective ARS ratings given by the participants (**Figure 5A**), it is not immediately clear whether ensemble modulation in PCG is more reflective of “bottom-up” information from the stimulus or the “top-down” ARS rating. Although, for many JD conditions, participants consistently gave almost all presentations of the stimulus the same ARS rating (e.g. T17 gave 98% of JD2 trials a “2”), for some JD conditions, the participants’ ARS rating wavered between two ratings (e.g. T11 rated 58% of JD1 trials a “3” and 39% a “2”). In the instances where there was a substantial split in ARS ratings for a given JD stimulus, there were no significant differences in ensemble FR between trials that were assigned different ARS ratings in response to the same stimulus. In contrast, neural responses (in 6d_T11_ and 6v_T17_) to different JD stimuli that were assigned the same ARS rating were found to be statistically different (RS tests, *p* < 0.05, **Figure 5C, Figure S7B**). This suggests that PCG activity was more closely reflective of the stimulus that was presented than of the participant’s ARS rating.

### Effect of Context - The “Aha!” Moment

Another variable that can affect (or effect) observation-related responses in motor cortex is the context under which stimuli are presented^21^. In the JD Task, participants understood that they would be viewing a set of stimuli of varying degrees of human-likeness and were instructed to provide ratings on how well they saw a hand (ARS rating). How might the motor cortex respond without this context?

In a separate session day prior to T11’s first JD Task session, we had the opportunity to capture neural activity in 6d_T11_ while T11 passively viewed dot stimuli *before* he was advised, and before he was aware, that they were meant to represent a hand (see “Naive Jittered Dots Task” in **SI Appendix**). We showed T11 four blocks of stimuli that were presented in increasing anthropomorphic order: (1) “Jittered” dot stimuli (similar to JD3), (2) “Semi-Jittered” dot stimuli (similar to JD2), (3) “Non-Jittered” dot stimuli (equivalent to JD0), and (4) Human Hand. Despite us giving him no instruction other than to passively observe the screen, during the first two blocks he made several comments expressing his curiosity in what he was viewing on the screen (e.g., “What are you trying to learn from this?”). During the middle of the 3rd block (after the 18th trial), T11 remarked, “Oh! I figured it out!” (see **Video S2**). After the fourth block, we asked him what he “figured out” in that moment, and T11 confirmed that he figured out that the dot stimuli he was viewing in Block 3 looked to him like a moving hand.

In congruence with what we would find in subsequent JD Task sessions (**Figure 2C**), the percentage of units responsive to the most hand-like dot stimulus (“Non-Jittered”) in Block 3 (up to 23%) was most similar to the percentage of units responding to the Human Hand stimulus in Block 4 (up to 24%, **Figure 6A**). However, the serendipitous remark made by T11 after trial 18 of Block 3, which we call his “Aha!” moment, also offered an opportunity to compare neural responses to the Non-Jittered dot stimulus before and after T11 was *aware* that the stimulus represented a hand. By evaluating ensemble activity trial-by-trial over the course of Block 3, a distinct and dramatic transition appeared around the time T11 said he “figured it out” after trial 18 of this block (**Figure 6B**). Specifically, we found that the ensemble FRs during observation of the Non-Jittered dot stimulus (JD0) *after* trial 18 were significantly higher than the trials *before* trial 18 (RS test, one-tailed, *p* < 0.01). Meanwhile, Empty observation trials did not exhibit significantly higher ensemble FRs after this “Aha!” moment (**Figure S8**), suggesting an increase in modulation specific to the JD stimuli and not a more general increase in neural activity over time. We also observed this effect on a single-unit level, wherein 19 units displayed increased FRs during Non-Jittered dot stimuli (Pinch trials, 12 units for Power trials) after the “Aha!” moment compared to before the “Aha!” moment, and only 5 units displayed increased FR responses to the Empty trials after the “Aha!” moment (**Figure 6C, Figure S8**).

**Figure 6.**
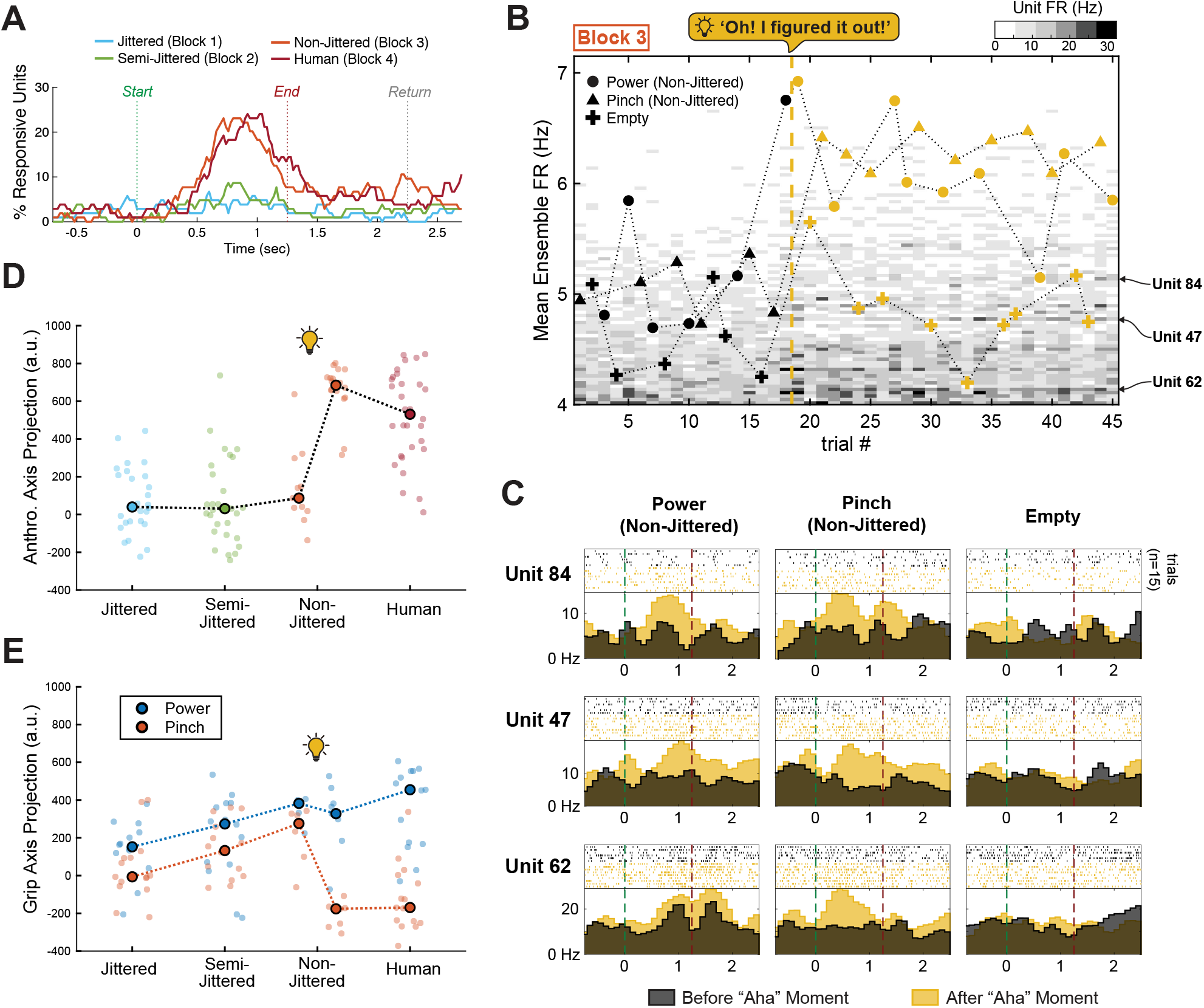
The “Aha!” Moment. (A) Percent of units responsive (see **SI Appendix**) when T11 viewed jittered dot stimuli for the first time. (B) Observed increase in neural modulation during Block 3 when T11 said that he “figured it out” that the moving dot animation represented a human hand. Grayscale image represents mean firing rates of each neuron during each trial during the block, where darker shading means higher firing rate (FR) during the trial. Units are ordered from most active (bottom) to least active (top). Overlayed traces denote mean ensemble FR for each trial. (C) Spiking activity during Block 3 of sample units that exhibited significantly increased FRs (RS test, *p* < 0.01) after the “Aha!” moment. Peristimulus time histograms were created using 100ms bins and smoothed with a 20ms Gaussian kernel. Mean FR changes for all recorded units are summarized in **Figure S8**. (D) Projections of the 10D SSIMS representations onto the anthropomorphicity “axis” for each stimulus condition in Blocks 1-4. Anthropomorphicity axis was defined here by the line connecting the SSIMS centroids of the Human (power and pinch) trials from Block 4 and the Empty trials from all blocks. Trials before (trials 1-17) and after (19-45) the “Aha!” moment are separated for emphasis. Connected dots represent medians for each group of trials. (E) Similar to (D), but showing the projections of the 10D SSIMS representations onto the grip “axis” for each stimulus condition. Grip axes for each session were defined by the line connecting the SSIMS centroids of all power and pinch trials (**SI Appendix**).

Neural ensemble spiking patterns were also affected by the “Aha!” moment. By projecting the 10D SSIMS representation of all trials onto an “anthropomorphicity axis” (see **SI Appendix**) we found that spiking activity during observation of the Non-Jittered stimuli became much more similar to that of the Human condition *after* the “Aha!” moment (**Figure 6D**). Likewise, by projecting the data onto a “grip axis” we found that the representations of power and pinch trials during Non-Jittered observation became much more distinct after T11’s realization (**Figure 6E**). Taken together, these results suggest that during observation of the Non-Jittered stimulus, the neural response became not only more robust, but also more similar to Human Hand observation after T11 made the cognitive association of the stimulus with a hand. This finding hints at the potential impact of context in the activity of motor cortex during action observation.

### Effects of Effector Anthropomorphicity During Attempted Movements

In order to enable voluntary control of an effector such as a robotic arm, iBCIs record and decode neural activity related to *attempted* or *imagined* movements of a participant. Therefore, to understand how effector anthropomorphicity may affect use of an iBCI, we evaluated neural activity in PCG while participants attempted hand grips while observing virtual effectors of different anthropomorphicities.

In the Virtual Effector (VE) Task (see **SI Appendix**), participants observed five “effectors” on a screen: the Human Hand condition (same as JD Task), an anthropomorphic Robot Hand, a Robot Claw, a Cube, and the Empty condition (**Figure 7A**). Effectors were presented pseudorandomly in an instructed delay paradigm wherein a Grip Cue – “power” or “pinch” – was audibly cued 1 - 2.5 seconds before a Go Cue (587 Hz tone) occurred corresponding with the start of a “power” or “pinch” movement performed by the effector (**Figure 7B, Video S3**). A baseline condition showing the Empty stimulus paired with a “relax” grip cue was also presented. Participants watched the virtual effectors under two contexts: Active trial blocks, in which they were instructed to *observe* and *attempt* the cued hand grips, and Passive trial blocks, in which they were instructed to only passively *observe* the effector movements on the screen (see **SI Appendix**). The same videos were viewed during Active and Passive trials. Participant T11 performed the VE Task across 3 session days and T17 performed one session of the VE Task.

**Figure 7.**
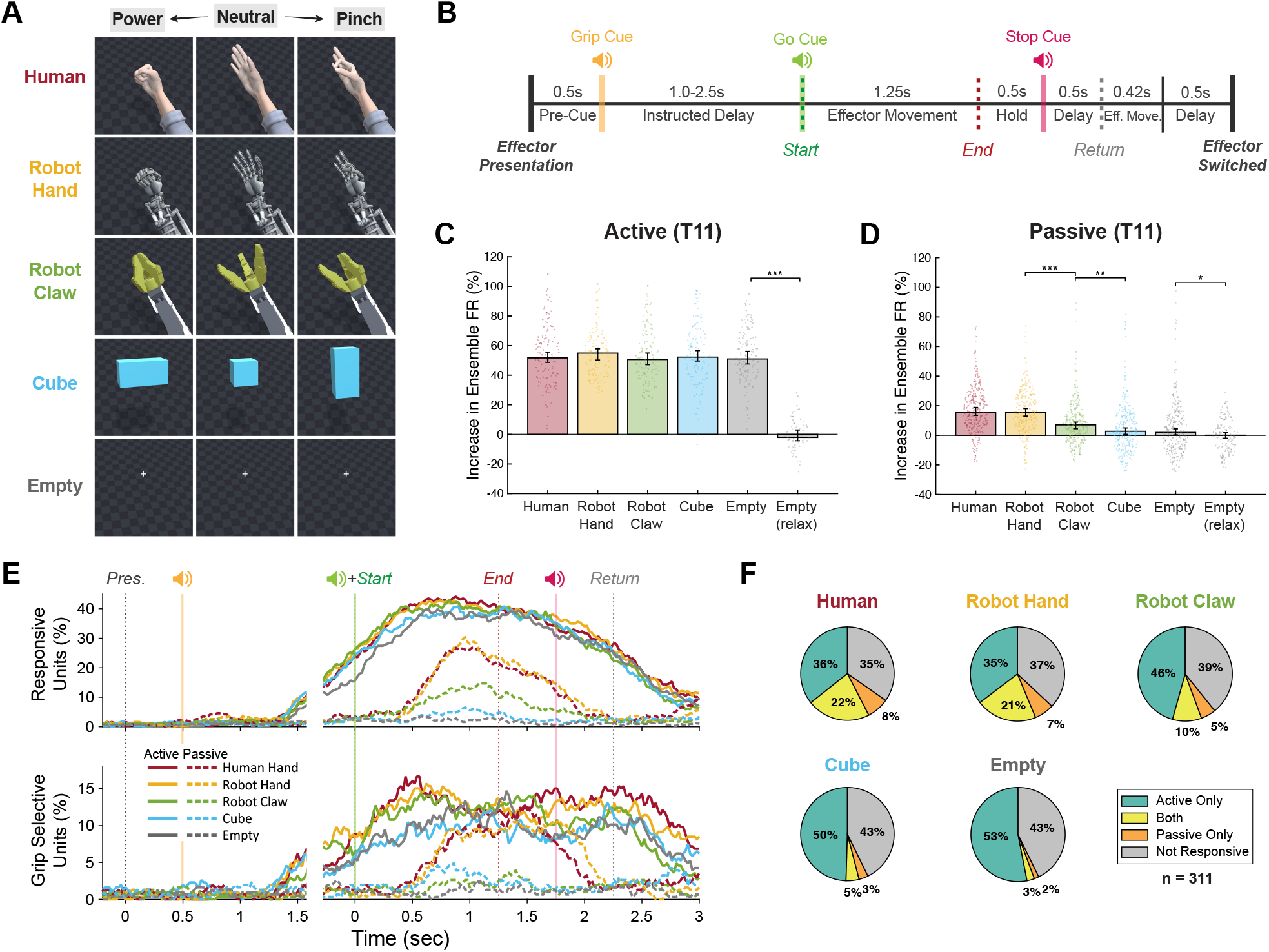
Comparison of Neural Responses During Active and Passive Performance of Virtual Effector Task. (A) Still frames from movies of the 5 virtual “effectors” shown to the participants. Effector postures at the start (“neutral”) and end (“power” or “pinch”) of the trial period are shown. See also **Video S3**. (B) Temporal structure of each trial. The grip cue (“power”, “pinch”, or “relax”), go cue, and stop cues were presented auditorily by the computer. See **SI Appendix** for task details. (C-D) Increases in 6d_T11_ FR during Active (C) and Passive (D) conditions while participant T11 watched each of the virtual effectors and the baseline condition (“Empty (relax)”). Mean ensemble FR increases are calculated and presented as in **Figure 2A-B** using data from the Empty-relax trials (Empty stimulus paired with a “relax” grip cue) as baseline. Significant decreases (Wilcoxon Rank Sum, one-tailed) between adjacent effector conditions are indicated with horizontal bars. Asterisks indicate significant *p*-values (*, *p* < 0.05; **, *p* < 0.01; ***, *p* < 0.001). (E) Percent of units in 6d_T11_ that were responsive during Active trials (solid lines) and Passive trials (dashed) over the course of the trial period for each effector type (see **SI Appendix**). (F) Percent of units in 6d_T11_ that were responsive during Active trials, Passive trials, or Both. Data from the time window 200ms-1700ms after the Go Cue was used to evaluate differences in FR between power, pinch, and baseline for each effector and motor intention condition (KW test, *p* <0.01). (G) Percent of units in 6d_T11_ that were selective for grip type. Units were considered grip selective in a given time window if they had significantly different (RS test, *p* < 0.01) FRs between power and pinch conditions.

#### Ensemble responses are similar across effector conditions during Active trials

As expected^3,5,22,23^, modulation of ensemble FRs in both participants was significantly greater during Active trials than during Passive trials (RS test, two-tailed: 6d_T11_, *p =* 2.48 × 10^-214^, 6d_T17_, *p =* 3.43 × 10^-125^, 6v_T17_, *p =* 3.08 × 10^-2^). However, Active trials did not produce significantly different modulation across the five effector conditions for either participant (**Figure 7C, Figure S9;** KW test: 6d_T11_, *p =* 0.83, 6d_T17_, *p =* 0.24, 6v_T17_, *p =* 0.70). Meanwhile, passive observation trials performed by T11 resulted in significant differences in 6d ensemble FR across effectors (KW, *p* = 2.82 × 10^-33^) that followed an anthropomorphic ordering (**Figure 7D**; Human, 15.6%; Robot Hand, 15.6%; Robot Claw, 7.0%; Cube, 2.7%; Empty, 1.9**%**). Despite robust observation-related neural activation in T17 during the JD Task (**Figure 2B**), neither ensemble recorded from T17 exhibited significant modulation during Passive trials of the VE Task (**Figure S9**). This may have been due to limitations in T17’s visual acuity at the time of VE Task data collection (see Discussion). Therefore, our findings from the VE Task will focus primarily on results from participant T11. (T17 results are also reported; **Figures S9, S10, S13**).

Similar to the ensemble FR, the proportions of responsive units (KW test, *p* < 0.01) during Active blocks were similar across effector conditions, with the highest percentage of responsive units occurring around 0.5 s after the Go Cue (**Figure 7E, Figure S10**). Neuronal activation also increased prior to the Go Cue, consistent with preparatory activity^24^. During Passive blocks, the proportion of responsive units in 6v_T11_ varied greatly, with the greatest proportion of neurons becoming active ∼1 second after the Go Cue during Human or Robot Hand trials (25.7% and 28.9%, respectively) and a much smaller proportion activating during Cube and Empty trials (6.4% and 3.5%, respectively). An intermediate number of responsive neurons were observed when the Robot Claw was passively observed, peaking at 14.5% of units 1.1s after the start of movement, reflecting a monotonic decrease in passive responses as the virtual effectors become less human-like (**Figure 7E**). Furthermore, when units were evaluated for *effector selectivity* (i.e. significant differences in FR across effectors, KW, *p* < 0.01), only up to 2.3% of 6d_T11_ units were effector selective during Active trials (**Figure S11**), suggesting that very few neurons presented significantly different responses to different observed effectors when hand grips were being attempted. By comparison, up to 43.7% of units were effector selective during Passive trials (**Figure S11**).

It was also informative to assess the overlap between neurons that responded during Active and Passive conditions (**Figure 7F, Figure S10**). In 6d_T11_, the proportion of neurons that were responsive (evaluated on 0.2-1.7s window after Go Cue) during both Active and Passive conditions (which some would characterize as exhibiting “mirror-like” properties) gradually increases when watching more anthropomorphic effectors **(Figure 7F)**. Notably, this increase is due to an overall increase in the number of neurons in 6d_T11_ responding to the Passive condition, whereas the overall number of neurons responding during the Active condition is relatively constant across effector types (∼55-58%).

#### Ensemble spiking patterns are similar across effector conditions during Active trials

A more precise understanding of the influence of anthropomorphicity on neural activity during Active and Passive trials can be gleaned by evaluating spike trains using SSIMS analysis (**Figure S12, Figure S13**). When spike trains patterns from 6d_T11_ are represented in a 2D similarity space, grip trials in the Active condition were well separated, demonstrating the expected markedly different ensemble firing patterns between power, pinch, and relax trials (**Figure S12A**). Although Passive trials were less separable by grip type, their SSIMS representations exhibited some clustering based on effector type, with Human and Robot Hand trials appearing more similar to Active power and pinch trials and Cube and Empty trials appearing more similar to Active relax trials (**Figure S12B**). Classification of the 10D SSIMS space confirmed that grips were easily separable for all effector types during Active trials (**Figure S12C**), with grip classification accuracies 89% or above that did not significantly differ across effectors (KW test, *p* = 0.29). Meanwhile, for Passive trials, grip classification accuracy differed by the observed effector (KW test, *p* = 4.78 × 10^-8^), with only the Human and Robot Hand conditions resulting in grip classification accuracies above chance (**Figure S12D**). Furthermore, neural classification of the *effector* during Active and Passive trials confirmed our conclusions from the 6d_T11_ population FR analysis: whereas neural activity patterns during Passive trials yielded above chance classification of the effector being observed, the effector type could not be identified via SSIMS classification from the neural activity that occurred during Active trials (**Figure S12E**). These findings indicate that despite significant anthropomorphic tuning in T11’s PCG during passive observation of movement, the impact of effector anthropomorphicity is greatly diminished when movement is simultaneously attempted.

## DISCUSSION

### Anthropomorphic Tuning in Human Premotor Cortex

In this study, intracortically recorded neuronal activity in the dominant precentral gyrus (PCG) of two people with tetraplegia was found to be tuned to the anthropomorphicity of observed movements. Specifically, passive observation of hand and hand-like animations demonstrated the presence of a gradient-like relationship between hand-likeness and neuronal ensemble activity in dorsal PCG (area 6d). Prior studies in humans and non-human primates have established that premotor cortex, especially its ventral portion, PMv, is highly associated with the observation of movements performed by others^7,25,26^ The robust neural responses in T11_6d_ and T17_6d_ (PMd) related to human hand observation are consistent with accounts of “mirror neurons” in monkey PMd^4,27^ and fMRI data suggesting mirror-like activation of human PMd^7^. However, the present study offers an unprecedented account of mirror-like activity in human premotor cortex and the dependence of this neural activity on an observed agent’s human-likeness.

The abundance of anthropomorphically tuned neural activity in PMd, reflected as a monotonic decrease in neural activation in area 6d ensembles as the observed effector becomes less human-like (**Figures 2, Figure 7**), can help explain conflicting results of neuroimaging studies that sought to determine whether robotic effectors elicited mirror-like activation in motor cortex^10–16^. As much as the subjective impression of a robot’s anthropomorphicity exists along a gradient^28^, so too does its neural representation in the cortical ensemble activity recorded in this study. The opportunity to record from individual neurons in the precentral gyrus allowed for a more spatially precise and direct exploration of neural responses than that available with PET, fMRI, or EEG. This allowed us to compare neural responses within sets of effectors that spanned a spectrum of anthropomorphic likeness. Because previous studies were limited to comparing the observation of human movement with only one or two robotic exemplars, the detected presence or absence of significant neural activation may have been dependent on the anthropomorphicity of the chosen stimuli. While the anthropomorphic gradient described for our two participants will benefit from replication in other subjects, our findings highlight the importance of exploring this effect and other gradient-like relationships throughout the sensorimotor system.

The anthropomorphic gradient we observed at the population level (**Figures 2-3**) was also reflected at the level of individual neurons. Single-unit recordings revealed a convincing preponderance of neurons that responded to the JD stimuli in accordance with their anthropomorphic ordering (**Figure 4**), suggesting that individual neurons themselves can be seen as “tuned” to anthropomorphicity. Furthermore, this suggests that population-level modulation of the motor cortex due to anthropomorphicity (**Figure 2A-B**) is a result of not only the gradual recruitment of additional neurons when watching increasingly hand-like effectors (**Figure 1B**) but also gradual increases in activity of individual neurons within the population. This result calls into question the value of discretely categorizing “mirror neurons” as distinct, specialized computational units in the brain. As prior critiques have noted^26,29^, the use of statistical testing to identify neurons that respond to both action production and action observation is prone to omitting weaker responses in neurons that do not rise above a particular threshold. Rather, our results suggest that it may be better to consider observation responses in the motor cortex as a network-level effect that variably modulates neurons depending on where they sit within the network. The outcome is a network-level encoding of sensory information that is comprised of neurons exhibiting a spectrum of levels of selectivity for movement observation. Aflalo et al^30^ drew similar conclusions based on the heterogeneity of mirror-like responses they observed among neurons in human posterior parietal cortex. Further work is needed to understand the single-unit and network representations of this anthropomorphic gradient as well as other continuous variables affecting observation responses in the numerous other brain regions that have been associated with mirror-like activity^7^. It is notable that observation responses from the neural ensemble in T17’s ventral premotor area, or area 6v (T17_6v_, **Figure S2**), were far less robust than responses from T17’s PMd (T17_6d_, **Figure 2**), considering PMv has historically been the most directly associated with mirror activity in motor cortex^26^. This may have been related to a more sparse sampling of single units that were isolated from T17’s 6v arrays during these sessions (**Table S2**). Alternatively, the limited responses of T17_6v_ during both Active and Passive conditions of the VE Task suggests the recorded ensemble may not be well tuned for the hand and finger movements assessed in this study. Future work that incorporates animations of more complex movements such as reaching and grasping objects^31^ may elicit more robust activation of 6v ensembles consistent with prior reports of visual responses in monkey and human PMv^2,7^.

Accurately quantifying the human-likeness of an observed stimulus is a notable challenge. Here, we defined and used the term “anthropomorphicity” to specifically evaluate the *visual similarity* of each effector with a human hand. Nevertheless, two effectors can be visually similar or different in several ways. For example, superficial variations such as coloring and the level of detail of a hand stimulus appear to not affect mirror responses in the motor cortex^32,33^, but robotic movement stimuli that significantly differ in their form and/or kinematics from human controls have been associated with reduced mirror activation in neuroimaging studies^10,11,13,14,34–37^. Our analysis of the neural activity in T11_6d_ during passive observation of the VE Task effectors appears to agree with these neuroimaging findings. The Human Hand and Robot Hand conditions, which differed superficially but were very similar in their forms and movement profiles, evoked similarly robust modulations of the ensemble and grip separability during passive observation (**Figure 7, Figure S12**). The Robot Claw, Cube, and Empty conditions, which more obviously differed from a human hand in their visual forms and movement, evoked distinctly muted responses.

Results of the JD Task indicate that varying either the form or the kinematics of an effector can independently modulate neural activity during observation. In the first three JD stimuli (JD0, JD1, JD2), the starting positions of the dots were increasingly randomized, while the individual trajectories of each dot during the movement phase of the trial were kept the same (see **SI Appendix, Video S1**). The discovery of a significant, and step-wise, reduction in neural activation as the starting positions of the dots became less human-like (**Figures 2-4**) lends support for the notion that the relative anthropomorphicity of an effector’s form is an important modulator of observation responses in PMd^10,11,14,34,38^. When we held the randomization of the starting points constant and increasingly varied movement trajectories (JD2-5), we also saw a small monotonic decrease in neural activation, which could be likened to fMRI studies that found differences in PMd activation when subjects observed human-like and non-human-like movements performed by the same agent^13,36^. Nevertheless, additional study is necessary to understand the relative importance of these variables and of even more nuanced differences in effector anthropomorphicity (e.g. robots with 3 fingers vs. 5 fingers, linear vs. parabolic velocity profiles, etc.) and how they interact with contextual inputs.

### Effects of Context on Anthropomorphic Tuning

Our primary objective in this study was to evaluate changes in neural activity in response to objective variations of a visual stimulus along an anthropomorphicity axis. However, the dramatic shift in neural activity corresponding with the moment T11 realized the dot stimuli represented a hand (**Figure 8**) demonstrates the powerful role higher-order cognitive influences may have on PMd activity during observation. Original accounts of “mirror” activity in the brain described visual activation of motor neurons as a largely innate, bottom-up process wherein understanding of other humans could arise from a “matching” of the incoming visual representation of an action with one’s own motor representation of the same action^39,40^. However, this “direct matching model” is challenged by later studies showing that observation responses in motor cortex can be modulated by a range of presumably “top-down” factors such as action context and prior experience^21^. A compelling demonstration of this effect has been fMRI studies showing that hemodynamic activity in premotor cortex can be visually evoked by simple geometric shapes after a learned association is made between the inanimate objects and human motor action^35,38,41^. Results of the Naive Jittered Dots Task suggest that the mental association of an inanimate visual stimulus with anthropomorphic movement can modulate both ensemble-level (**Figure 6B,D-E**) and individual neuron (**Figure 6C, Figure S8**) responses in PMd. Furthermore, the temporal specificity of intracortical recordings combined with T11’s “Aha!” moment provided rare evidence that this cognitively driven shift in PMd excitability during observation can occur over the course of only a few seconds.

Press^37 49^ proposed that the biological tuning of mirror responses in the brain is the result of years of learning to associate observed movements performed by other humans with executed (or imitated) movements performed by one’s self and that reduced activation of mirror neuron areas during robot observation could be explained by a weaker *association* of the stimulus with a familiar sensorimotor experience. Our results (**Figure 6**) suggest that activity in PMd during observation is influenced by a *conscious* (at least implicitly recognized) association between the stimulus and anthropomorphic movement. After this anthropomorphic association with the JD stimuli was established (i.e. in the sessions following the “Aha!” moment; **Figures 2-5**), neural activity appeared to more directly reflect subtle variations in the visual stimuli rather than the participant’s subjective (or conscious) assessment of anthropomorphicity (**Figure 5C**). While the neural activity in PMd was associated with the participant’s assigned ARS score (**Figure 5B**), that same neural activity was insufficient to generate the mental conclusion about the similarity or quality of the subtly different stimuli (**Figure 5C**). It will be interesting to explore whether this suggests that a ‘vote’ from PMd could be overridden by other mental processes, or that this PMd activity is not involved in the processes that result in the expressed conscious perception of the stimuli.

These results imply significant contributions of both “top-down” knowledge and “bottom-up” visual information in the neural responses of PMd during observation. Hypotheses for a functional purpose of mirror phenomena in motor cortex would benefit from incorporating both of these information sources in their models. Kilner et al.^42^ proposed one such model wherein mirror responses result from the involvement of the motor system in a predictive coding scheme that compares top-down movement predictions with bottom up sensory inputs^42–44^. One interpretation of our results could be that the anthropomorphic gradient we found in PMd was driven by the relative utility of the human motor system in modeling *predictions* of upcoming movements in these anthropomorphically ambiguous effectors (see Supplementary Discussion 1). Nevertheless, aside from our conclusion that both top-down and bottom-up factors likely affect (or effect) observation responses in human PMd, a more definitive understanding of this complex interaction and their potential functional importance is left to future study.

### Effects of Effector Type during Attempted Movement

A goal of intracortical BCI research is to provide an intuitive, information-rich input for volitional control of effectors including assistive robotics, prosthetic limbs, or functional electrical stimulation-supported limb reanimation. Over more than two decades of iBCI research, people with tetraplegia have used hand and arm imagery to control a wide range of devices from simple computer cursors^45,46^ to high-dimensional anthropomorphic robots^47–49^. One motivation for this study was to evaluate whether the anthropomorphicity of an iBCI effector can meaningfully affect neural activity in the motor cortex and thereby affect the decoding of intent.

The VE Task demonstrated how attempted hand grasps (similar to the motor imagery one might use to control an iBCI) substantially impact neural activity in area 6d. For T11, passive observation of the VE Task effectors yielded differences in neural activation of 6d (**Figure 7, S12**) consistent with the anthropomorphic gradient revealed by his neural responses in the JD Task (**Figure 2-3**). However, during “Active” trials of the VE Task, differences across effector conditions were heavily attenuated (**Figure 7, S12**). Although the results from Active trials with T17 support the conclusion of a minimized impact of visual feedback in 6d during attempted grasping (**Figure S9**), the nearly absent neural responses observed during Passive trials, which were not consistent with earlier JD Task results, made it such that we could not rule out other factors affecting these results. For example, for several weeks following performance of the JD Task (Trial Day 62), T17 reported decreased visual acuity as a consequence of exposure keratitis related to weakness of his orbicularis oculi muscles. With treatment, T17 reported improvement in his visual acuity by the time the VE Task was performed (Trial Day 90), however it is possible T17’s vision was still moderately affected on this day. Furthermore, whereas the JD Task incorporated a participant feedback period in its trial design (“ARS rating”, **Figure 1C**), the VE Task did not include a behavioral component to confirm visual acuity or attention. We were unable to repeat the VE Task with T17, but future repetitions of the VE Task would benefit from including an ARS feedback period as a behavioral control and to facilitate comparisons with the JD Task.

Our finding that grip classification accuracies (power vs. pinch) during the Active condition (**Figure S12C**) did not significantly differ between effectors suggests that the anthropomorphicity of a chosen actively-controlled effector may not significantly affect iBCI performance. However, it should be noted that this is a relatively simple classification task for modern iBCI systems, and it is unclear whether subtle changes due to effector appearance could have a greater impact on higher dimensional decoding tasks (see also Supplementary Discussion 2).

Our results suggest that in this “open-loop” setting, the attempted movement signal during the Active condition may have obscured the prominent anthropomorphic tuning we observed in area 6d during the Passive condition (in the absence of attempted movement). It will be interesting to study whether effector anthropomorphicity has a greater impact on neural activity in the motor cortex during “closed-loop” control where visual feedback from the effector is more directly relevant to task performance and could have short-latency impacts on subsequent neural activity. Further studies that evaluate the impacts of effector anthropomorphicity during closed-loop iBCI control would not only improve our understanding of sensorimotor integration in the brain, but also help inform the optimal design of robotic effectors for use by people with iBCIs.

## METHODS

This study reports on results from 7 research sessions performed with participant T11 and 2 research sessions with participant T17, who each gave informed consent and are enrolled in a pilot clinical trial of the BrainGate2 Neural Interface System. Enrollment criteria and details about the trial can be found at http://www.clinicaltrials.gov/ct2/show/NCT00912041 (Caution: Investigational device. Limited by Federal law to investigational use). Permissions for this study were granted by the US Food and Drug Administration (FDA, Investigational Device Exemption #G090003) and the Institutional Review Boards of Massachusetts General Hospital (#2009P000505), VA Providence Healthcare, and Brown University. Detailed methods for neural recording and spike-sorting, MRI reconstructions, task details (JD Task, Naive JD Task, VE Task), and statistical analyses are provided in the **SI Appendix**.

## ACKNOWLEDGEMENTS

We thank T11, T17, and their care partners for their dedicated contributions to this research; B. Travers, D. Rosler, and M. Masood for administrative support; Z. Williams for surgical planning and implanting of T11’s and T17’s arrays; S. Mernoff for his role as site-responsible clinical investigator; J. Simeral for engineering support; and N. Schmansky and S. Cash for their help producing the MRI-derived brain reconstructions. This work was supported by the American Heart Association (19CSLOI34780000), Office of Research and Development, Rehabilitation R&D Service, Dept of Veterans Affairs (N2864C, A2295R), NIH NINDS (UH2NS095548), NIH NIDCD (U01DC017844), the Movement Disorder Foundation, the ALS Association, and the Cerebral Palsy Alliance Research Foundation. The funders had no role in study design, data collection and interpretation, or the decision to submit the work for publication.

CAUTION: Investigational Device. Limited by Federal Law to Investigational Use. The content is solely the responsibility of the authors and does not necessarily represent the official views of the National Institutes of Health, or the Department of Veterans Affairs, or the United States Government.

## Notes

https://github.com/jgusman/anthropomorphicity

https://doi.org/10.5061/dryad.37pvmcw08

### Competing Interest Statement

Z.C.B. is currently an employee at Google LLC, Mountain View, CA. T.S.S-C is currently with the Biomedical Engineering Graduate Group, University of California Davis, Davis, CA. A.K. is currently with the Drexel University College of Medicine, Philadelphia, PA. T.H. is currently with the Department of Bioengineering, University of California Berkeley, Berkeley, CA and the Department of Bioengineering and Therapeutic Sciences, University of California San Francisco, San Francisco, CA. J.P.D is a paid Director and shareholder of Pathmaker Neurosystems and an advisor and shareholder for Neurable and Beacon Biosignals. None of these entities provided support for this project. The MGH Translational Research Center has a clinical research support agreement (CRSA) with Ability Neurotech, Axoft, Neuralink, Neurobionics, Paradromics, Reach Neuro, Precision Neuro, and Synchron, for which LRH provides consultative input. Mass General Brigham (MGB) convenes the Implantable Brain-Computer Interface Collaborative Community (iBCI-CC); charitable gift agreements to MGB, including those received to date from Axoft, Blackrock Neurotech, Neuralink, Paradromics, Precision Neuro, and Synchron, support the iBCI-CC, for which LRH provides effort.

## REFERENCES

1. di Pellegrino, G., Fadiga, L., Fogassi, L., Gallese, V. & Rizzolatti, G. Understanding motor events: a neurophysiological study. Exp. Brain Res. 91, 176–180 (1992).

2. Gallese, V., Fadiga, L., Fogassi, L. & Rizzolatti, G. Action recognition in the premotor cortex. Brain 119 (Pt 2), 593–609 (1996).

3. Tkach, D., Reimer, J. & Hatsopoulos, N. G. Congruent activity during action and action observation in motor cortex. J. Neurosci. 27, 13241–13250 (2007).

4. Papadourakis, V. & Raos, V. Neurons in the macaque dorsal premotor cortex respond to execution and observation of actions. Cereb. Cortex 29, 4223–4237 (2019).

5. Dushanova, J. & Donoghue, J. Neurons in primary motor cortex engaged during action observation. Eur. J. Neurosci. 31, 386–398 (2010).

6. Vigneswaran, G., Philipp, R., Lemon, R. N. & Kraskov, A. M1 corticospinal mirror neurons and their role in movement suppression during action observation. Curr. Biol. 23, 236–243 (2013).

7. Molenberghs, P., Cunnington, R. & Mattingley, J. B. Brain regions with mirror properties: a meta-analysis of 125 human fMRI studies. Neurosci. Biobehav. Rev. 36, 341–349 (2012).

8. Maeda, F., Mazziotta, J. & Iacoboni, M. Transcranial magnetic stimulation studies of the human mirror neuron system. Int. Congr. Ser. 1232, 889–894 (2002).

9. Caspers, S., Zilles, K., Laird, A. R. & Eickhoff, S. B. ALE meta-analysis of action observation and imitation in the human brain. Neuroimage 50, 1148–1167 (2010).

10. Tai, Y. F., Scherfler, C., Brooks, D. J., Sawamoto, N. & Castiello, U. The human premotor cortex is ‘mirror’ only for biological actions. Curr. Biol. 14, 117–120 (2004).

11. Perani, D. et al. Different brain correlates for watching real and virtual hand actions. Neuroimage 14, 749–758 (2001).

12. Kuz, S. et al. Mirror neurons and human-robot interaction in assembly cells. Procedia Manufacturing 3, 402–408 (2015).

13. Cross, E. S. et al. Robotic movement preferentially engages the action observation network. Hum. Brain Mapp. 33, 2238–2254 (2012).

14. Gazzola, V., Rizzolatti, G., Wicker, B. & Keysers, C. The anthropomorphic brain: The mirror neuron system responds to human and robotic actions. Neuroimage (2007) doi:10.1016/j.neuroimage.2007.02.003.

15. Kupferberg, A. et al. Fronto-parietal coding of goal-directed actions performed by artificial agents. Hum. Brain Mapp. 39, 1145–1162 (2018).

16. Oberman, L. M., McCleery, J. P., Ramachandran, V. S. & Pineda, J. A. EEG evidence for mirror neuron activity during the observation of human and robot actions: Toward an analysis of the human qualities of interactive robots. Neurocomputing 70, 2194–2203 (2007).

17. Phillips, E., Zhao, X., Ullman, D. & Malle, B. F. What is human-like?: Decomposing robots’ human-like appearance using the anthropomorphic roBOT (ABOT) database. in Proceedings of the 2018 ACM/IEEE International Conference on Human-Robot Interaction (ACM, New York, NY, USA, 2018). doi:10.1145/3171221.3171268.

18. Crowell, C. R., Deska, J. C., Villano, M., Zenk, J. & Roddy, J. T., Jr. Anthropomorphism of robots: Study of appearance and agency. JMIR Hum. Factors 6, e12629 (2019).

19. Epley, N., Waytz, A. & Cacioppo, J. T. On seeing human: a three-factor theory of anthropomorphism. Psychol. Rev. 114, 864–886 (2007).

20. Victor, J. D. & Purpura, K. P. Nature and precision of temporal coding in visual cortex: a metric-space analysis. J. Neurophysiol. 76, 1310–1326 (1996).

21. Kemmerer, D. What modulates the Mirror Neuron System during action observation?: Multiple factors involving the action, the actor, the observer, the relationship between actor and observer, and the context. Prog. Neurobiol. 205, 102128 (2021).

22. Vargas-Irwin, C. E. et al. Watch, Imagine, Attempt: Motor cortex single-unit activity reveals context-dependent movement encoding in humans with tetraplegia. Front. Hum. Neurosci. 12, 450 (2018).

23. Rastogi, A. et al. Neural representation of observed, imagined, and attempted grasping force in motor cortex of individuals with chronic tetraplegia. Sci. Rep. 10, 1429 (2020).

24. Crammond, D. J. & Kalaska, J. F. Prior information in motor and premotor cortex: activity during the delay period and effect on premovement activity. J. Neurophysiol. 84, 986–1005 (2000).

25. Rizzolatti, G., Fadiga, L., Gallese, V. & Fogassi, L. Premotor cortex and the recognition of motor actions. Brain Res. Cogn. Brain Res. 3, 131–141 (1996).

26. Kilner, J. M. & Lemon, R. N. What we know currently about mirror neurons. Curr. Biol. 23, R1057–62 (2013).

27. Fogassi, L. et al. Visual responses in the dorsal premotor area F2 of the macaque monkey. Exp. Brain Res. 128, 194–199 (1999).

28. Mori, M., MacDorman, K. & Kageki, N. The uncanny valley [from the field]. IEEE Robot. Autom. Mag. 19, 98–100 (2012).

29. Heyes, C. & Catmur, C. What happened to mirror neurons? Perspect. Psychol. Sci. 17, 153–168 (2022).

30. Aflalo, T. et al. Cognition through internal models: Mirror neurons as one manifestation of a broader mechanism. bioRxiv 2022.09.06.506071 (2022) doi:10.1101/2022.09.06.506071.

31. Vargas-Irwin, C. E., Franquemont, L., Black, M. J. & Donoghue, J. P. Linking Objects to Actions: Encoding of Target Object and Grasping Strategy in Primate Ventral Premotor Cortex. Journal of Neuroscience 35, 10888–10897 (2015).

32. Désy, M.-C. & Lepage, J.-F. Skin color has no impact on motor resonance: evidence from mu rhythm suppression and imitation. Neurosci. Res. 77, 58–63 (2013).

33. Zhu, H., Sun, Y. & Wang, F. Electroencephalogram evidence for the activation of human mirror neuron system during the observation of intransitive shadow and line drawing actions. Neural Regeneration Res. 8, 251–257 (2013).

34. Biermann-Ruben, K. et al. Right hemisphere contributions to imitation tasks. Eur. J. Neurosci. 27, 1843–1855 (2008).

35. Engel, A., Burke, M., Fiehler, K., Bien, S. & Rösler, F. What activates the human mirror neuron system during observation of artificial movements: bottom-up visual features or top-down intentions? Neuropsychologia 46, 2033–2042 (2008).

36. Casile, A. et al. Neuronal encoding of human kinematic invariants during action observation. Cereb. Cortex 20, 1647–1655 (2010).

37. Press, Clare. Action observation and robotic agents: Learning and anthropomorphism. Neurosci. Biobehav. Rev. 35, 1410–1418 (2011).

38. Engel, A., Burke, M., Fiehler, K., Bien, S. & Rosler, F. How moving objects become animated: the human mirror neuron system assimilates non-biological movement patterns. Soc. Neurosci. 3, 368–387 (2008).

39. Gallese, V. & Goldman, A. Mirror neurons and the simulation theory of mind-reading. Trends Cogn. Sci. 2, 493–501 (1998).

40. Rizzolatti, G., Fogassi, L. & Gallese, V. Neurophysiological mechanisms underlying the understanding and imitation of action. Nat. Rev. Neurosci. 2, 661–670 (2001).

41. Clare Press et al. fMRI evidence of ‘mirror’ responses to geometric shapes. PLoS One 7, e51934 (2012).

42. Kilner, J. M., Friston, K. J. & Frith, C. D. Predictive coding: an account of the mirror neuron system. Cogn. Process. 8, 159–166 (2007).

43. Kilner, J. M. More than one pathway to action understanding. Trends Cogn. Sci. 15, 352–357 (2011).

44. Friston, K., Mattout, J. & Kilner, J. Action understanding and active inference. Biol. Cybern. 104, 137–160 (2011).

45. Hochberg, L. R. et al. Neuronal ensemble control of prosthetic devices by a human with tetraplegia. Nature 442, 164–171 (2006).

46. Pandarinath, C. et al. Inferring single-trial neural population dynamics using sequential auto-encoders. Nat. Methods 15, 805–815 (2018).

47. Hochberg, L. R. et al. Reach and grasp by people with tetraplegia using a neurally controlled robotic arm. Nature 485, 372–375 (2012).

48. Collinger, J. L. et al. High-performance neuroprosthetic control by an individual with tetraplegia. Lancet 381, 557–564 (2013).

49. Wodlinger, B. et al. Ten-dimensional anthropomorphic arm control in a human brain-machine interface: difficulties, solutions, and limitations. J. Neural Eng. 12, 016011 (2015).

