## Supplementary material for "Observation-Related Activity in Human Motor Cortex Increases with Effector Anthropomorphicity": SI Appendix

### DETAILED METHODS

#### Study Permissions and Participant Details

This study reports on results from 7 research sessions performed with participant T11 and 2 research sessions with participant T17, who each gave informed consent and are enrolled in a pilot clinical trial of the BrainGate2 Neural Interface System. Enrollment criteria and details about the trial can be found at http://www.clinicaltrials.gov/ct2/show/NCT00912041 (Caution: Investigational device. Limited by Federal law to investigational use). Permissions for this study were granted by the US Food and Drug Administration (FDA, Investigational Device Exemption #G090003) and the Institutional Review Boards of Massachusetts General Hospital (#2009P000505), Providence VA Medical Center, and Brown University.

T11 is an ambidextrous man, 36-years-old at the time of the study, who has tetraplegia due to a C4 AIS-B spinal cord injury that occurred 11 years prior to enrollment in the trial. He has some maintained proximal arm strength, but little or no voluntary movement at his wrists and fingers bilaterally. T17 is a right-handed man, 33-years-old at the time of the study, who is completely paralyzed and anarthric due to advanced ALS (ALSFRS-R = 0). He uses eye movements to communicate with his care partners and medical staff.

Participant T11 had two 96-channel NeuroPort microelectrode arrays (4mm x 4mm, 1.5mm electrode length, platinum tips; Blackrock Neurotech, Salt Lake City, UT) surgically implanted in the "hand knob" area of the left precentral gyrus, which was identified by pre-operative magnetic resonance imaging (MRI). Participant T17 had six 64-channel NeuroPort microelectrode arrays (3.2mm x 3.2mm, 1.5mm electrode length, iridium oxide) implanted in motor- and speech-related areas along the left precentral gyrus. **Figure 1A** shows array placement locations registered to MRI-derived reconstructions of the participants' brains.

#### 3D Brain Reconstruction and Electrode Array Localization

MRI-derived reconstructions of the participants' brains (**Figure 1A**) were generated using FreeSurfer (https://surfer.nmr.mgh.harvard.edu) and rendered in Blender (https://www.blender.org) using MMVT (https://github.com/pelednoam/mmvt). To identify electrode locations relative to the reconstructed cortex, we performed a spatial co-registration of the pre-implantation MRI scan with a post-implantation CT scan and confirmed placement by comparing to photographs of the implanted electrodes in the open craniotomy. The shaded areas in **Figure 1A** represent regions defined by the Human Connectome Project (HCP) HCP-MMP1.0 atlas[1], which were generated using an HCP annotation mapped to FreeSurfer space[71] and a set of FreeSurfer commands (https://github.com/ftadel/IntrAnat/issues/7) that converted the labels to native space such that they could be mapped to the 3D model in Blender. The resultant output (**Figure 1A**) showed both of T11's electrode arrays and the two most medial of T17's arrays in *Area 6d* of the HCP parcellation, the two most lateral of T17's arrays in *Area 6v*, and the two remaining of T17's arrays in *Area 55b*. To confirm these observations, we regenerated the HCP parcellations by processing the preoperative MRI T1 and T2 scans through the Minimal Processing Pipeline[2] and using Connectome Workbench (https://www.humanconnectome.org/software/connectome-workbench) commands to map the HCP parcellation back to native space. The latter approach showed a similar mapping of array locations.

#### Neural Recording and Spike-Sorting

Neural signals from T11's electrode arrays were analog filtered between 1 – 7.8k Hz, digitally sampled at 20 kHz, and transmitted wirelessly using two 'Brown Wireless' devices (Blackrock Neurotech, Salt Lake City, UT) and wall-mounted antennae. The 20 kHz data was upsampled to 30 Hz sampling rate (via sample-and-hold) for compatibility with downstream processing. This wireless recording setup was shown previously to have negligible differences in signal quality compared to the wired recording setup[3]. Neural signals from T17's electrode arrays were analog filtered between 0.3 – 7.5 kHz and digitally sampled to 30 kHz using two Neuroplex E (Blackrock Neurotech) devices and transmitted via cables. Custom software based in MATLAB and Simulink (MathWorks, Natick, MA) was developed to store and align up to 256-channels of 30 kHz neural data to trial events. Because of this channel count limitation, for sessions with participant T17, we chose to record from the four 64-channel arrays that were determined to modulate most with attempted hand-movements: two arrays from 6d and two from 6v.

Single-unit spiking activity was isolated offline by identifying differences in spike waveform shapes and amplitudes using custom-made software[4]. Units were then verified manually using Offline Sorter (Plexon Inc., Dallas, TX). Across the seven recording sessions with T11, a median of 96 units and 12 units were isolated from his medial 6d array and lateral 6d array, respectively, per session (see **Tables S1-S2** for the total number of units included in analysis for each session for each participant). Neural data recorded from T11's lateral array during Session 2 were discovered following the session to have been corrupted during saving to disk and thus not included in this analysis. However, because of the relatively few units isolated from the lateral array in other sessions, we decided to still include Session 2 in our analysis using only data from the medial array. Note that the groups of neurons recorded from day to day using microelectrode arrays can potentially overlap. Therefore, the sets of isolated units from each recording session should be considered as partially overlapping samples of the total pool of cortical neurons.

#### Jittered Dots Task

Participant T11 performed the Jittered Dots (JD) Task on post implant trial days 385, 387, and 412. Participant T17 performed the JD Task on post implant day 62. During this task, we recorded neural activity during the passive observation of a simple anthropomorphic dot animation (Unity Technologies, San Francisco, CA) that was systematically altered to appear progressively less like a human hand. In addition to the 6 dot stimulus conditions (referred to as Jittered Dots 0 (JD0), Jittered Dots 1 (JD1), etc.), we used the Human Hand and Empty conditions from the Virtual Effector Task (see below) to represent the "most anthropomorphic" and "least anthropomorphic" limits, respectively, of our *anthropomorphicity* spectrum (**Figure 1A**, **Figure S1**, **Video S1**). JD0 was generated by simply taking the 3D model of the Human Hand, removing the "skin" of the model, and aligning 26 white, spherical dots to the underlying joints of the model. As a result, JD0 represents the same kinematics as the original Human Hand grasp animations but without the superficial appearance of the Human Hand model. In JD1 and JD2, the starting positions of the dots were randomly shifted relative to JD0 while maintaining the same three-dimensional trajectories of the dots used in JD0. In JD3 and JD4, the randomization of the starting positions of the dots was left the same as JD2, but the movement trajectories of each dot were rotated in an increasingly random way (see **Table S3** for details). JD5 represented the most "random" condition where the dots moved in linear translations in random directions unrelated to the kinematics of the original hand model. Note that JD2-5 were visually identical up until the start of effector movement. All effectors were shown performing distinct "power" and "pinch" grip movements, except for the JD5 and Empty conditions, which showed the same type of animation for both grip conditions (**Figure S1**).

All blocks in the JD Task were Passive blocks; participants were instructed to fix gaze on the center of the screen while keeping their body relaxed (see *Virtual Effector Task* section below for more on Active vs. Passive blocks). In addition, participants were instructed to describe their subjective assessment of the anthropomorphicity of each presented stimulus at the end of each trial using the Anthropomorphicity Rating Scale (ARS) defined as follows:

"4" = I definitely saw a hand
"3" = I think I saw a hand
"2" = I think I did NOT see a hand
"1" = I definitely did NOT see a hand

After each trial, T11 would verbally say the number corresponding to his evaluation of the prior trial and a researcher would record his response. Because T17 is unable to speak, to indicate his ARS rating after each trial, he would make saccadic eye movements to numeric (1-4) labels positioned at the left, top, right, and bottom edges of the computer monitor showing task stimuli. One researcher with prior experience reading T17's eye movements for communication would call out the chosen ARS rating and another researcher would write it down.

The inclusion of the ARS was done for two reasons: to confirm that our *a priori* assumption of the relative anthropomorphicity of the JD stimuli matched the participant's subjective experience and to confirm that the participant maintained attention throughout the task. Trials in which the participant did not give a response after a trial were removed from analysis.

JD Task trials were structured identically to those of the VE Task (see below) but included an additional period (0.75s for T11, 3.0s for T17) at the end of each trial where a green circle would be shown and the participant would communicate their ARS rating **Figure 1B**). Unlike the VE Task, no grasp cues were used, thus the participant could not "anticipate" the upcoming grip movement prior to trial start. Furthermore, not including Active blocks during JD Task sessions helped decrease the possibility that the participant might accidentally "attempt" movement during blocks he was supposed to perform passively. Each effector was shown 6 times (3 times per grip) per block, and 13-16 blocks were run each session (**Tables S1-S2**).

#### Naive Jittered Dots Task

Prior to running the JD Task with T11, we devoted one session (Session 4, Trial Day 372) to capturing his neural responses to the dot stimuli before and after he had recognized the association of the stimulus with a human hand. The plan for this session was to start by having T11 passively watch several blocks of jittered dot stimuli without telling him that the "visual stimuli" were related to hands. Then, unless he spontaneously recognized the hand-like stimuli, we would explicitly tell him that the stimuli were supposed to appear hand-like and have him watch more blocks of the same stimuli to see whether his neural responses changed due to the new contextual information. Additionally, to ensure the participant did not make the anthropomorphic association too early, we structured the "before" blocks as follows:

Block 1 - Jittered (a highly jittered dot condition, similar to JD2)
Block 2 - Semi-Jittered (a "less" jittered dot condition, similar to JD1)
Block 3 - Non-Jittered (non-jittered, equivalent to JD0)
Block 4 - Human Hand

Each block displayed 45 trials total: 15 "power" grip trials, 15 "pinch" grip trials, and 15 Empty trials (with a white fixation cross), which were presented in a randomized order.

Unfortunately, due to an error in the experimental code that was discovered after the session, we realized that the set of stimuli that were shown after we explicitly confirmed the contextual information was slightly different than the set of stimuli shown prior to informing him. Thus, we determined that direct comparisons between neural responses to the "before" blocks and the "after" blocks would not be meaningful to report. However, the spontaneous remark offered by T11 during Block 3 (see Results, **Figure 6**) provided a demarcation that allowed us to evaluate changes in neural responses between sets of trials before and after this remark (or, "Aha Moment"). Note that this particular novel stimulus presentation could not be repeated with the same participant, since, for T11, the dot stimuli would be associated with a hand for the foreseeable future). This unique finding is presented as inspiration for future studies to explore with additional participants.

**Virtual Effector Task**

Participant T11 performed the Virtual Effector (VE) Task on post implant trial days 262, 276, and 280. Participant T17 performed the VE Task on post implant day 90. With this task, we explored neural responses while participants viewed five different effector types (**Figure 7A**, **Video S3**): a Human Hand, an anthropomorphic Robot Hand, a Robot Claw with three "fingers", a blue Cube, and the Empty condition, where a static white fixation cross was shown. Using Unity (Unity Technologies, San Francisco, CA), each effector (except the Empty condition) was animated to produce two types of movements corresponding to two common grip conditions: "power" and "pinch". The Human Hand and Robot Hand were animated to produce anatomically accurate "power" and "pinch" grip movements, as shown in **Figure 7A**. The Robot Claw represented a "power" grip by flexing all three of its fingers and represented a "pinch" grip by flexing two fingers. The Cube, which was designed to represent an "effector" that moved in a clearly non-anthropomorphic manner, was animated to increase its width during a "power" grip and increase its height during a "pinch" grip. In the Empty condition, which was designed as a baseline condition, no movement occurred in either "power" or "pinch" conditions.

Each stimulus was presented using an instructed delay paradigm (**Figure 7B**). The effector was presented in a neutral starting position in the center of the screen, before a grasp cue (e.g. "power") was communicated audibly by the computer. After a variable delay, a "beep" (Go Cue, 587 Hz tone) occurred, coinciding with the start of effector movement toward its target posture. The effector was "held" in the target posture for 0.5s before another "beep" (Stop Cue, 523 Hz) occurred, there was another 0.5s delay, and then the effector reversed its movement (at 3x speed) back to the neutral position (see **Video S3**). Due to an inadvertent difference in the stimulus presentation software, for T11, all "power" stimuli (i.e. VE Task and JD Task stimuli) began movement 375ms after the Go Cue and "pinch" stimuli began movement at the Go Cue. Both "power" and "pinch" movements were completed 1.25s after the Go Cue.

Therefore, T11 viewed power grip animations that were slightly delayed and slightly faster than the pinch grip trials. For T17, all animations (power and pinch) started and ended movement as expected at the Go Cue and 1.25s after the Go Cue, respectively. During the Pre-Cue epoch, the Instructed Delay epoch, and the final Delay epoch, the effector was shown in its unmoving Neutral position (**Figure 7A**). At the end of each trial, the current effector was immediately exchanged for the next effector. In total, there were 11 trial conditions for this task: a "power" and "pinch" trial for each of the effector types plus one additional Empty trial which was paired with a "relax" grasp cue. Trials were presented in blocks of 44 pseudo-randomized trials (4 of each stimulus condition).

Each VE Task session consisted of 2 types of blocks: *Active* blocks and *Passive* blocks (counts shown in **Tables S1-S2**). Both block types presented identical auditory and visual feedback but differed in the instructions given to participants at the start of each block. Before *Active* blocks, participants were given the following instructions:

> *"In this block during the trial we want you to fixate your eyes on the objects on the screen, and during the time between the Go Cue and the Stop Cue we want you to ATTEMPT and hold the requested hand grip. Make sure to continue attempting the hand movement until you hear the Stop Cue. And after the Stop Cue you may relax until the next grip cue."*

During *Passive* blocks, the instructions were as follows:

> *"During this block, you will see the same thing, but this is a PASSIVE task. Meaning your only instruction is to look at the objects on the screen. It is important that you DO NOT attempt the grasps that you are shown. It is also important that you try to stay focused on the hand and do not look away from the screen."*

Prior to running each block type for the first time, a practice block was run in order to ensure the participant understood the instructions for the given block type. Note that audio cues were present in both Active and Passive blocks of the VE Task, even though the participant did not need them to complete the Passive trials. No motor decoding feedback was provided to the participant during either trial type (i.e., trials were all "open loop").

During all sessions, T11 sat in his wheelchair, which supported his head, back, and legs. His arms were relaxed with his hands lying in a prone position on his lap. Small movements of T11's right hand were observed when he attempted hand grasps during Active blocks but not during Passive blocks, which served as additional confirmation that he was following the instructions for a particular block. The monitor was placed at a comfortable distance for T11's viewing, between 91-99 cm from his face. During all sessions with T17, he was lying in bed with his head and torso raised to view the monitor which was placed between 56-64 cm from his face.

#### QUANTIFICATION AND STATISTICAL ANALYSIS

All analyses were performed using MATLAB 2021b.

##### Statistical Analysis of Neural Firing Rates

Population and single-unit firing rates were evaluated across grip and effector conditions for each task (**Figure 2A-B**, **Figure 4**, **Figure 4B-C**, **Figure 7C-D, Figure S2A, Figure S7, Figure S9**). For each unit, mean trial firing rates were calculated on 1.5s windows of data starting 200ms after the start of effector movement. The "ensemble firing rate" represents the average taken over all isolated single units that were recorded in a given trial. In order to account for potential nonstationarities within and across session days, we represented modulation of the ensemble as the relative increase of the ensemble firing rate as a percentage of the block baseline firing rate. Block baselines were calculated as the mean ensemble firing rate of all Empty/Relax trials in a given block (in the JD Task this refers to all trials where the Empty condition is shown, whereas in the VE Task this refers to only trials where the Empty condition is shown paired with a "Relax" cue).

In order to evaluate the significance of our observation that ensemble firing rates decreased with decreasing anthropomorphicity, we performed one-tailed Wilcoxon Rank Sum tests ($p < 0.05$) between stimulus conditions. In the JD Task, we were interested in understanding whether neural modulation *gradually* decreased with decreasing anthropomorphicity or if the relationship was more of a binary, step-like function (high anthropomorphicity → high response, low anthropomorphicity → low response). We did not know *a priori* where each of our JD stimuli would fall along the theorized anthropomorphicity spectrum, thus it was conceivable that we could miss a graduated, though relatively steep transition in the neural response if our conditions were too widely or irregularly spaced. However, we reasoned that if there existed at least one condition that produced a distinctly intermediate neural response in motor cortex, this was evidence that the relationship between anthropomorphicity and neural modulation was *gradual* rather than binary. Thus, we used multiple Wilcoxon Rank Sum tests to test whether there existed multiple conditions that produced a neural response that was significantly *lower* than at least one *more* anthropomorphic condition and significantly *greater* than at least one *less* anthropomorphic condition. Furthermore, where possible, we presented the data from each condition as a swarm plot of individual trials to confirm that these intermediate responses represented unimodal distributions and were not the result of an averaging of bimodally high and low responses.

##### "Responsive" and "Selective" Units

Another way that we evaluated the response of the neural ensemble to each condition was by calculating the proportion of recorded single units that were "responsive" during a given condition and that were "selective" for a given condition subtype. Units were considered responsive if they displayed significantly different firing rates between the power grip variant of a stimulus, the pinch grip variant of that stimulus, and the baseline (Empty/Relax) condition (Kruskal-Wallis test, $p < 0.01$). Units were considered grip selective if their firing rates differed significantly between the power and pinch variants of a given stimulus. Likewise, effector selective units (**Figure S11**) demonstrated significant differences in firing rates between effector conditions for a given grip category. Note that by these definitions, all grip selective units are necessarily responsive as well. To calculate responsive and selective units over time (**Figures 2C-D**, **Figure 6A**, **Figures 7F-G, Figures S2B-C, Figures S10A,C,D,F**), we first collapsed spiking activity from each unit into 20ms bins of firing *rates* and smoothed the result with a 120ms gaussian kernel. Then for each unit we compared the distributions of responses to each condition as time-averaged firing rates within a 300ms sliding window that was shifted in 20ms steps across the trial period. When presented as a pie chart (**Figure 7F**, **Figures S10C,E**), unit responsiveness was evaluated on a single time window 200ms-1700ms after the Go Cue.

**Ensemble spike train similarity space analysis (SSIMS)**

SSIMS is an unsupervised dimensionality reduction algorithm designed to capture the intrinsic relationships between neural firing patterns on a trial-by-trial basis[5]. The algorithm starts by calculating pairwise distances between spiking patterns for each trial of each single cell using the metrics developed by Victor & Purpura[6]. This method utilizes the concept of "edit distances" where one spike train is transformed into another using a series of operations: adding a spike, deleting a spike, or moving a spike in time. The first two operations are assigned a "cost" equal to 1, and moving a spike in time is assigned a cost equal to $q\Delta t$, where $\Delta t$ represents the duration of the time shift and q is a parameter that determines the temporal precision of the metric. In our analyses, q was chosen so that moving a spike by more than 100 ms was equivalent to deleting the spike and then inserting another spike. The next step of the SSIMS algorithm uses t-SNE[7] to reduce the dimensionality of the concatenated pairwise distance matrices for each neuron in the ensemble, resulting in a map that preserves local neighborhood structure. In our implementation of the algorithm, t-SNE was initialized using principal component analysis (PCA). Each point on the SSIMS map corresponds to a trial, and the distance between the trials indicates the degree of similarity between the corresponding spiking patterns.

Using SSIMS, we generated low-dimensional projections of the spike train population data for each session to visualize the relative similarities between neural data resulting from each condition. To quantify these relationships, we created discrete classifiers for grip type and stimulus condition using the SSIMS data. For each session, neural activity patterns of all the sorted units were projected onto a 10-dimensional SSIMS representation generated using data from a 1.5s window starting 200ms after the start of effector movement. We used k-nearest neighbor (KNN) classification with 5 neighbors and 5-fold cross validation to generate decoding accuracies that can be interpreted as describing the relative differences between neural ensemble activity during each condition. Chance distributions for each figure were generated by repeating the above procedure using randomly shuffled trial labels 100 times for each dataset and combining across datasets in a given figure. Note that here "dataset" means the subset of trials used to generate each classification accuracy. For example, in **Figure 3C** there are 24 datasets corresponding to the 8 effector conditions and 3 sessions; thus the dashed lines represent the 95% confidence interval of the combined distribution of 2,400 "chance" accuracy calculations. When three sessions of a task were available (i.e. for T11), we assessed the significance of differences in decoding accuracies using the Kruskal-Wallis test (for **Figures S12C-D**) and pairwise Wilcoxon Rank Sum tests (**Figures 3C,F**, **Figure 12E**). Here, we treated each cross validation "fold" as an individual sample, allowing for 15 classification accuracies for each condition.

Although the SSIMS algorithm evaluates pairwise comparisons between individual trial spike trains in an unsupervised manner, structural patterns in the dataset can become apparent when viewing labeled low-dimensional projections of the data. In our 2D plots of the data from the JD Task (e.g. **Figure 3A**) we observed that data points tended to fall along an "axis" corresponding with the anthropomorphic arrangement of the stimuli conditions. To confirm this, we calculated the centroids (mean coordinate location over a set of trials) of the 10D SSIMS data from all the Human and Empty trials in a given session and defined a one-dimensional line that would represent this "axis". Then we projected each data point onto this line such that we acquired its relative spike train similarity to the "average" Human and "average" Empty trial (**Figure 3B, Figure S3B**, **Figure S4B**) . One benefit of this approach is that it allows us to combine SSIMS data across sessions by aligning the data to two anchor points that are common across sessions and restricting our analysis to the relative distances between those anchor points.

We performed a similar transformation using centroids defined by the set of all power and all pinch trials to create a "grip axis" (**Figure 3D**). We plotted the results for power and pinch trials separately for each condition to emphasize the gradual separation of the power and pinch neural representations as the participant observed increasingly anthropomorphic stimuli (**Figure 3E**, **Figure S3E**, **Figure S4E**).

#### ADDITIONAL RESOURCES

ClinicalTrials.gov Identifier: NCT00912041, registered June 3, 2009.

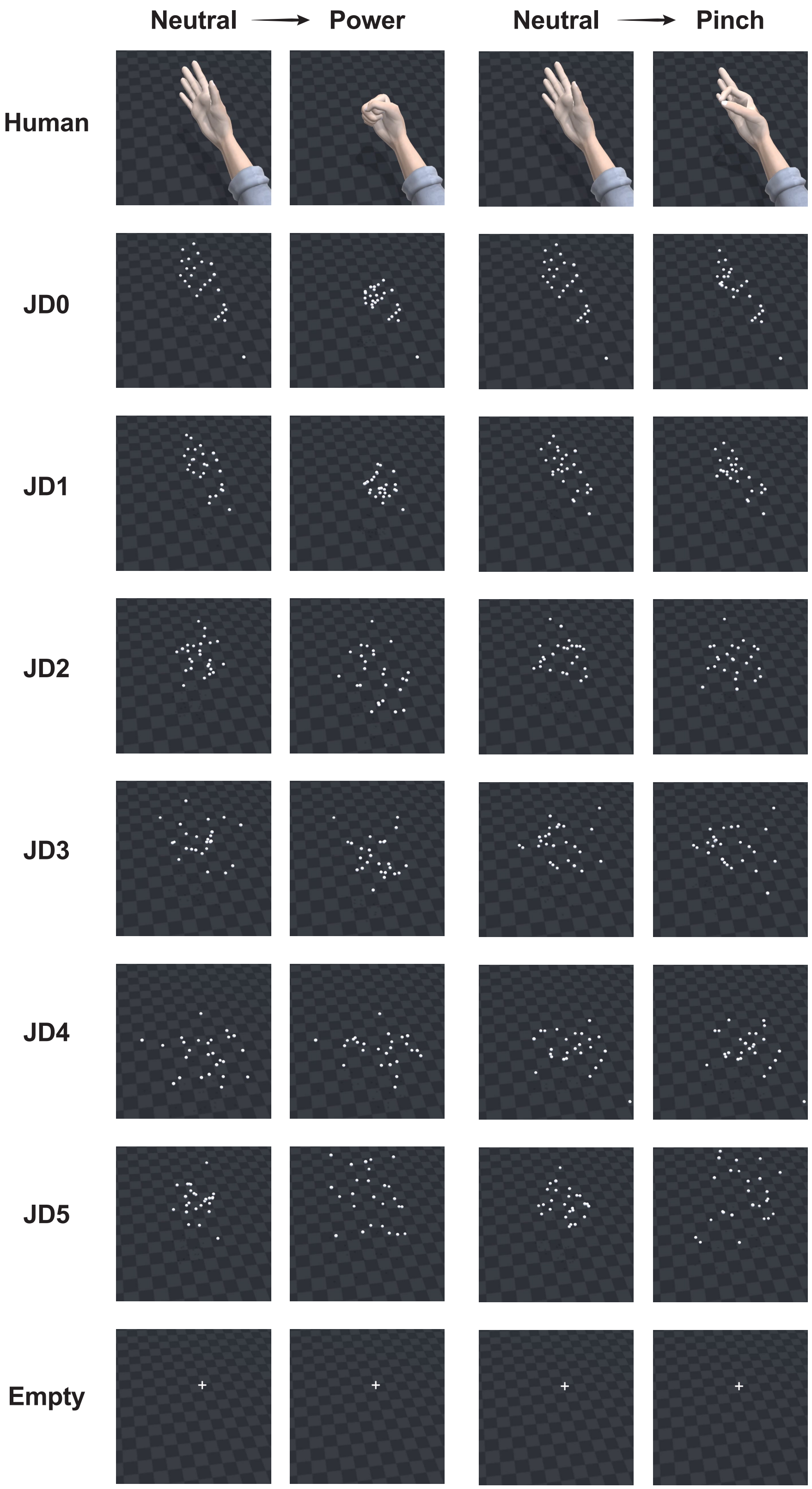


**Figure S1. Still Frames from the Beginning and End of All Stimulus Conditions Shown During the JD Task, Related to Figure 1.**

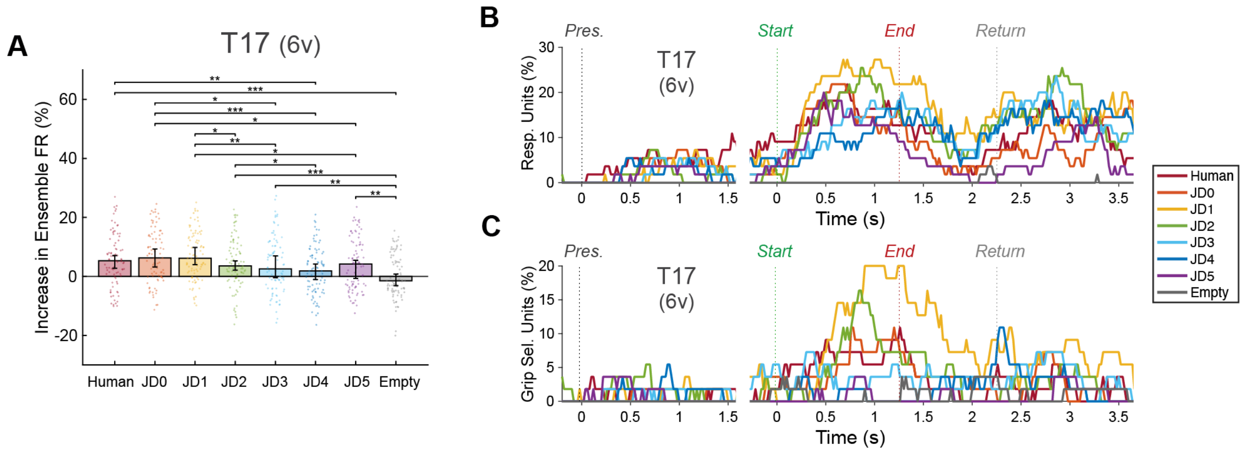


**Figure S2. Modulation of the Motor Cortical Ensemble During the Jittered Dots Task (T17 6v results). Related to Figure 2**

(A) Increases in ensemble firing rate (FR) in T17 area 6v during passive observation of each of the eight stimuli presented during the JD Task. See Figure 2 legend for details.
(B) Percent of units responsive to the task over the course of the trial period, plotted as in Figure 2C.
(C) Percent of units selective for grip type, plotted as in Figure 2D.

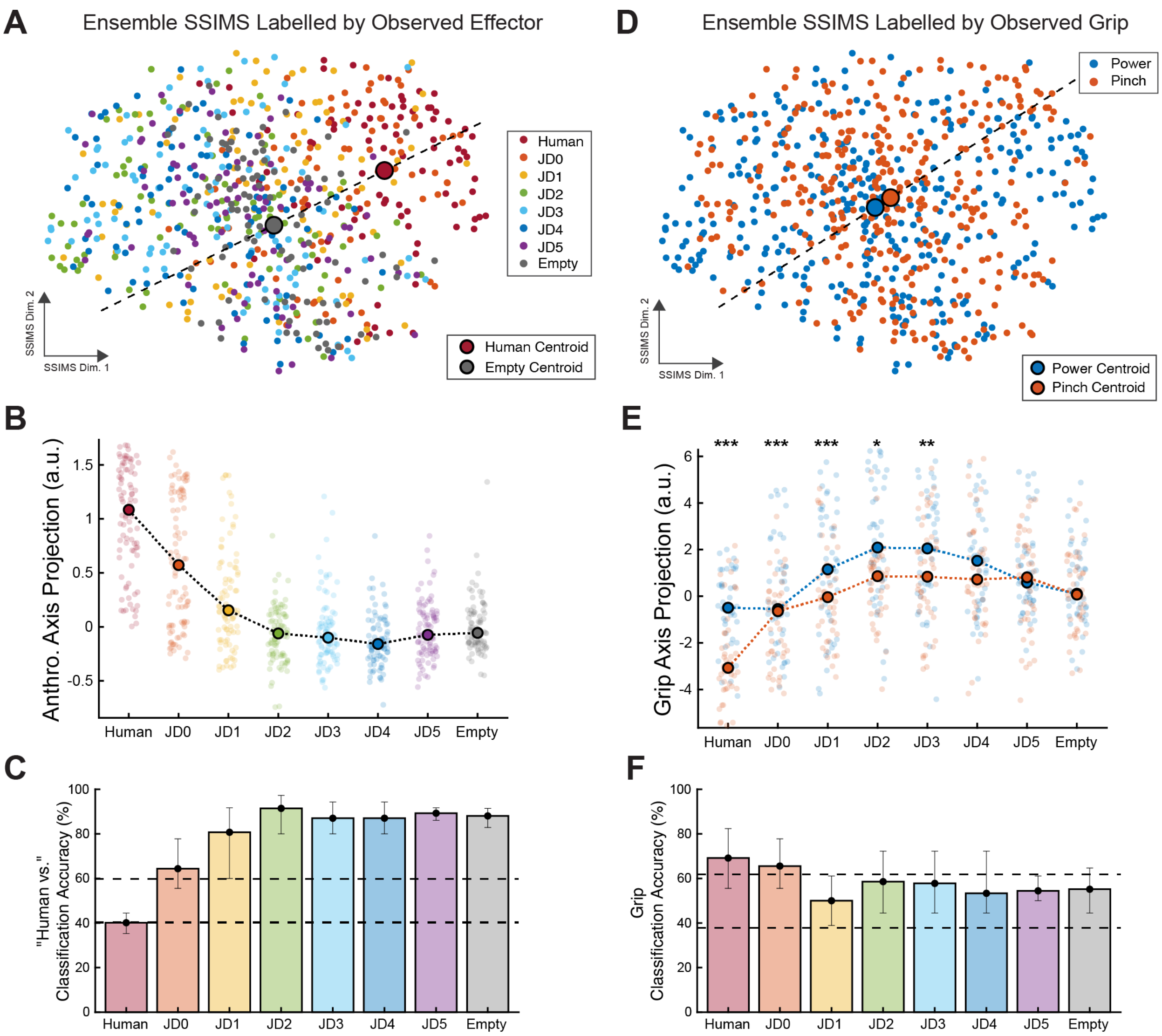


**Figure S3. Spike Train Similarity Space Analysis on Ensemble Firing Patterns in Area 6d of T17 during JD Task Session, Related to Figure 3**

See Figure 3 legend for details.

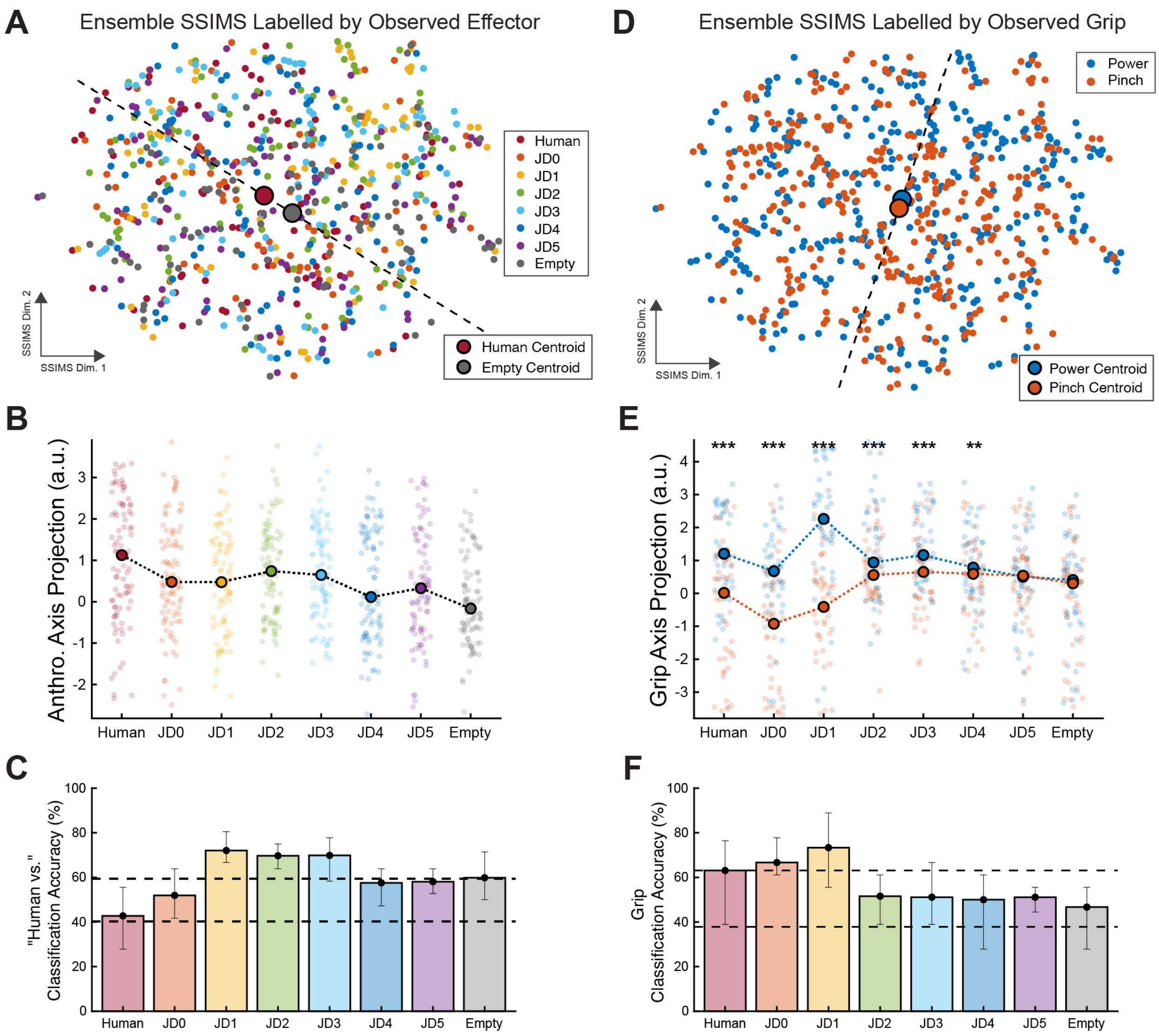


**Figure S4. Spike Train Similarity Space Analysis on Ensemble Firing Patterns in Area 6v of T17 during JD Task Session, Related to Figure 3**

See Figure 3 legend for details.

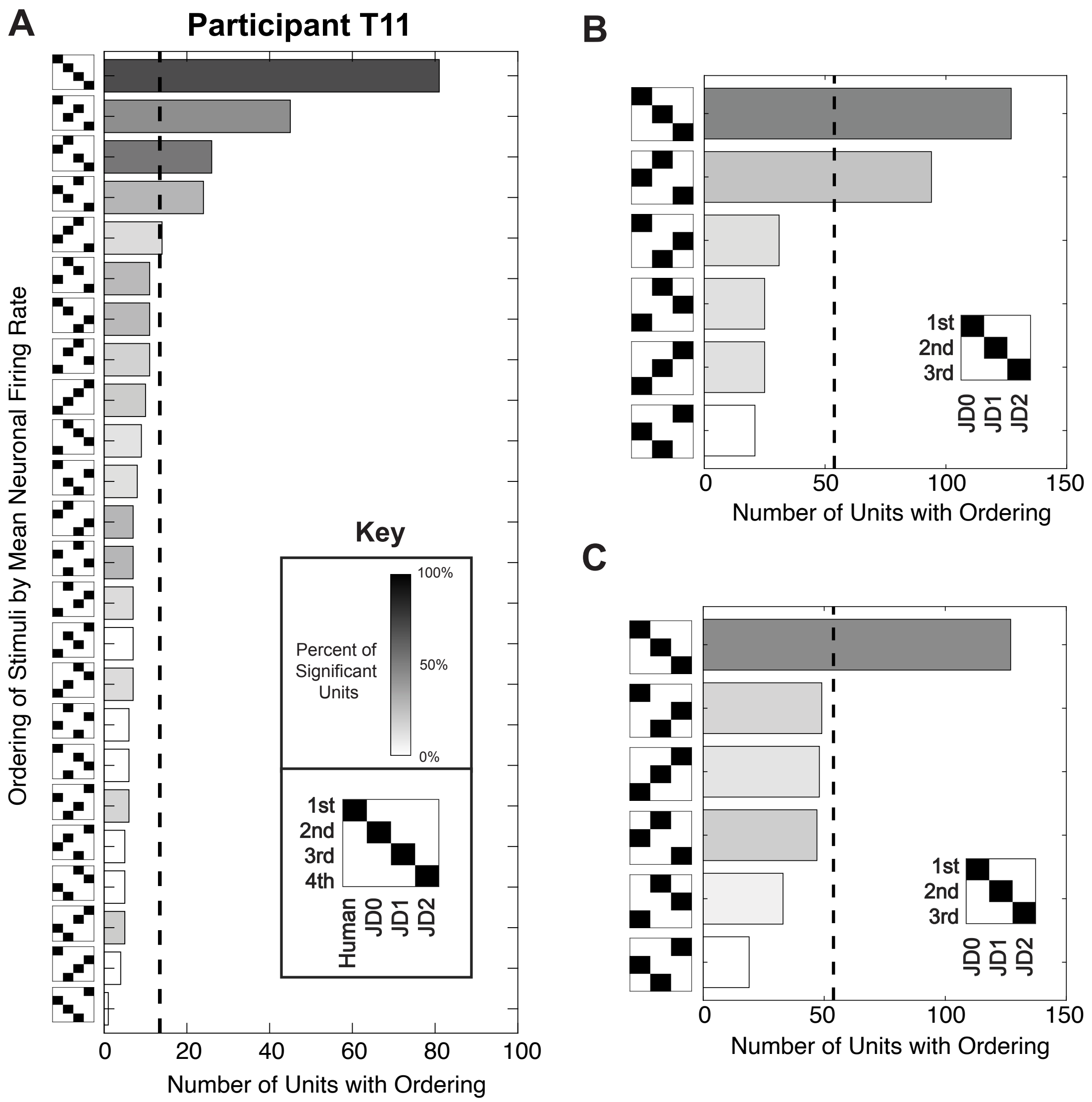


**Figure S5. Additional Analysis on Single-Unit Firing Rate Orderings in Participant T11, Related to Figure 4.**

(A) Neurons categorized by their relative responses to Human, JD0, JD1, and JD2 stimuli (“pinch” trials only). For each unit, responses to each stimulus condition were ranked according to the resultant mean firing rate during the 200-1700ms window following the start of stimulus movement. Bar plot lengths represent the number of neurons that had a particular ordering of mean firing rates. All 24 possible orderings are shown. The vertical dashed line reflects the number of neurons that would have a particular ordering by chance (~13.5). The color (grayscale) of each bar represents the percent of units within a category that have significantly different firing rates between two or more conditions (Kruskal-Wallis, $p < 0.01$).

(B) Same as (A), but neurons are categorized by their relative responses to JD0, JD1, and JD2 only. Number of neurons that would have a particular ordering by chance is 53.8.

(C) Same as (B), but with power trials.

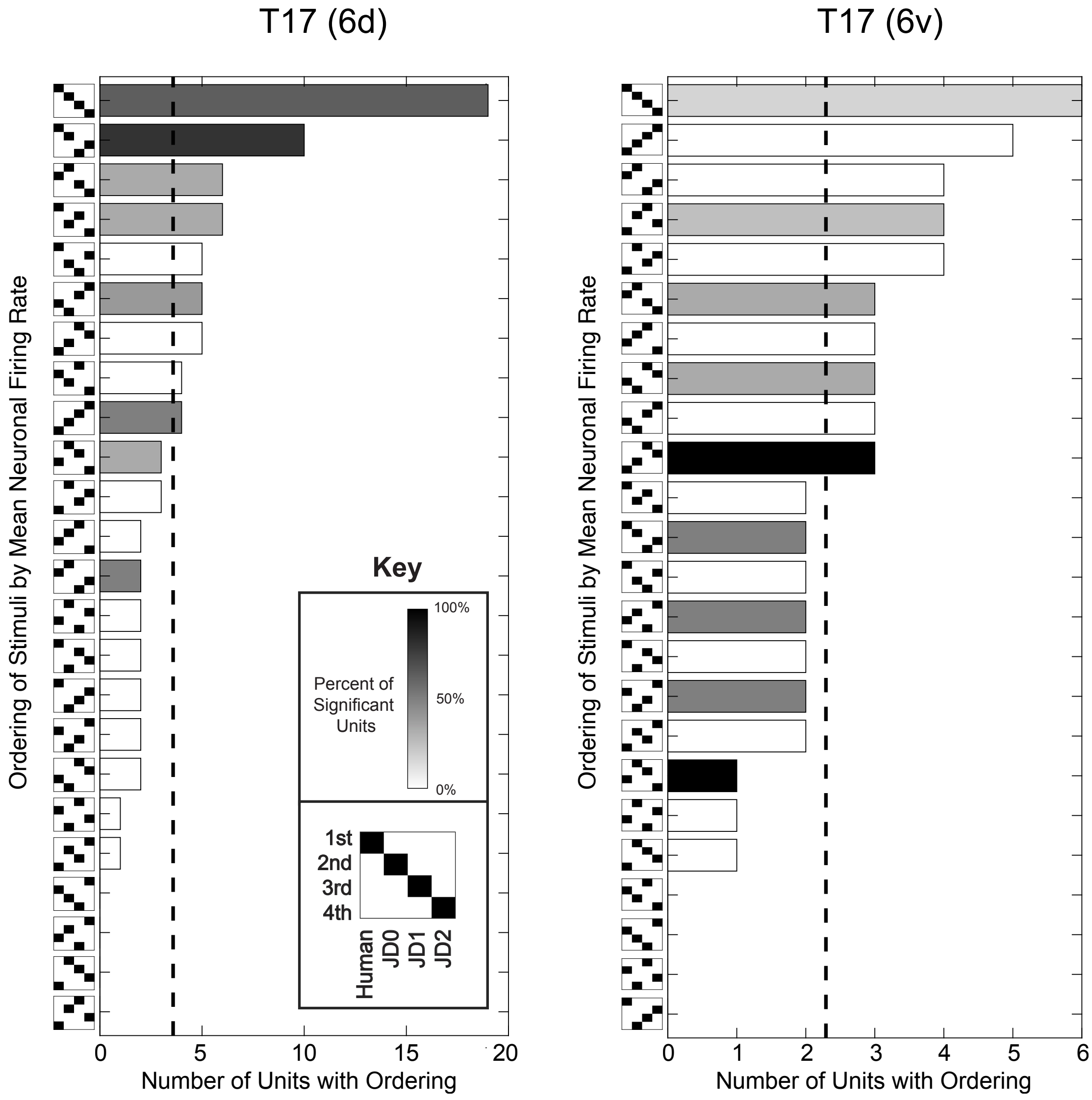


**Figure S6. Analysis of Single-Unit Firing Rates in Participant T17, Related to Figure 4**

Neurons from area 6d (left) and area 6v (right) in T17 categorized by their relative responses to Human, JD0, JD1, and JD2 stimuli ("power" trials only). For each unit, responses to each stimulus condition were ranked according to the resultant mean firing rate during the 200-1700ms window following the start of stimulus movement. Bar plot lengths represent the number of neurons that had a particular ordering of mean firing rates. All 24 possible orderings are shown. The vertical dashed line reflects the number of neurons that would have a particular ordering by chance (~13.5). The color (grayscale) of each bar represents the percent of units within a category that have significantly different firing rates between two or more conditions (Kruskal-Wallis, $p < 0.01$).

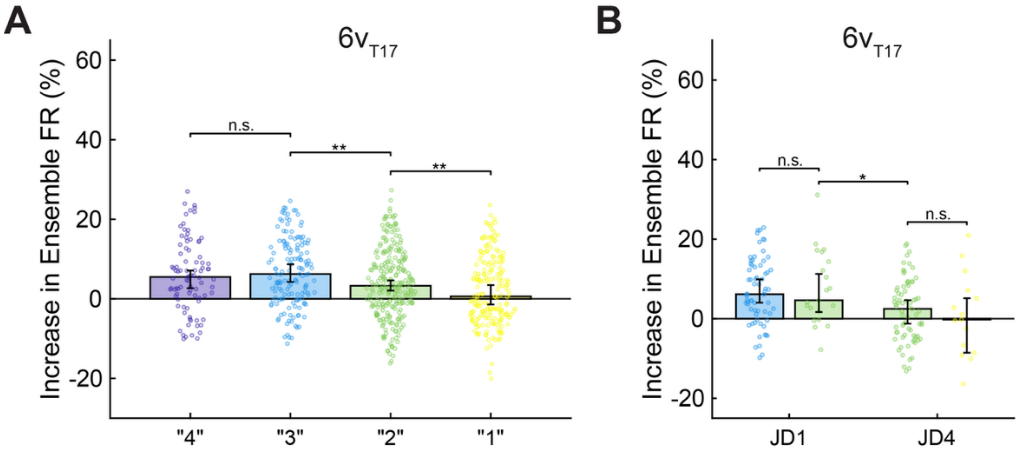


**Figure S7. Effects of ARS Rating on Ensemble Firing Rate in Participant T17 Area 6v, Related to Figure 5**

(A) Increases in 6v ensemble firing rate (FR) for all JD Task trials performed by T17, organized by the ARS rating given by the participant after each trial. Swarm plots, bar plots, error bars, and significant differences are presented as in **Figure 5B**.

(B) Increases in $6v_{T17}$ ensemble FR of trials for each stimulus condition, according to the given ARS rating. Only stimulus conditions that had more than 15 trials of more than one rating are shown. Swarm plots, bar plots, error bars, and significant differences are presented as in **Figure 5C**. No significant differences were found between trials of the same stimulus but different ARS rating, but in $6v_{T17}$, a significant difference ($p = 0.027$) in ensemble FR was found between trials of JD1 and trials of JD4 that were given "2" ARS ratings.

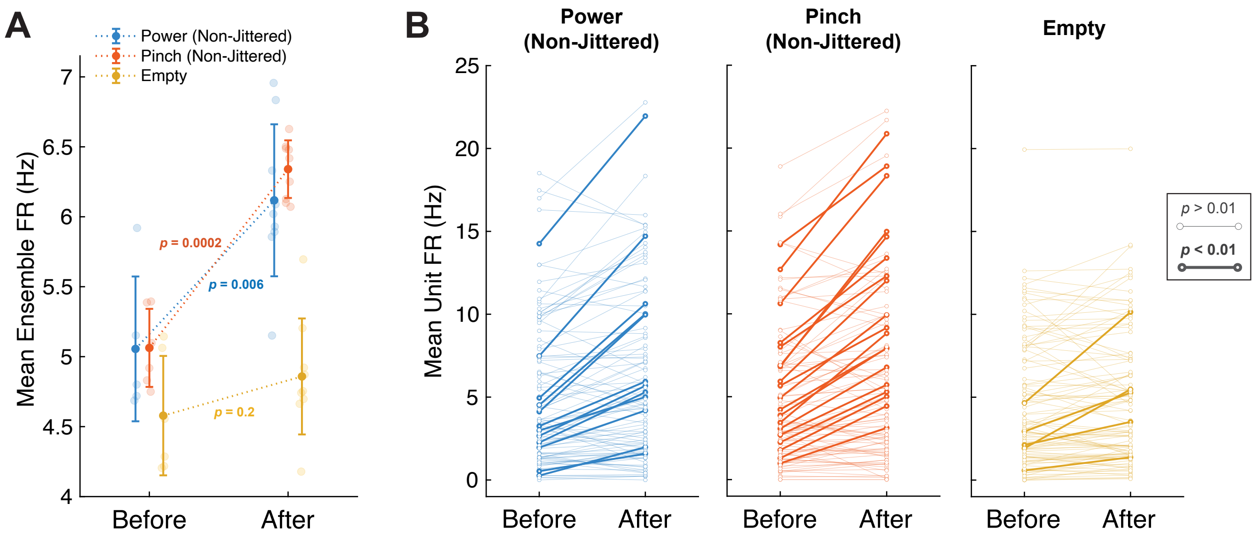


**Figure S8. Significant Ensemble and Single-Unit Firing Rate Changes Before and After the "Aha!" Moment, Related to Figure 6.**

(A) Ensemble FRs before (trials 1-17) and after (19-45) the "Aha!" moment for each type of stimulus in Block 4. Error bars denote means and standard deviations. The neural ensemble responses to the Non-Jittered dot stimulus increased significantly (RS test, $p < 0.01$, one-tailed) after the "Aha!" moment, whereas there was no significant increase in ensemble FR between Empty trials before and after the "Aha!" moment ($p > 0.01$).

(B) Mean single-unit FRs before and after the "Aha!" moment. Statistically significant increases (RS test, $p < 0.01$, one-tailed) are denoted by bold lines.

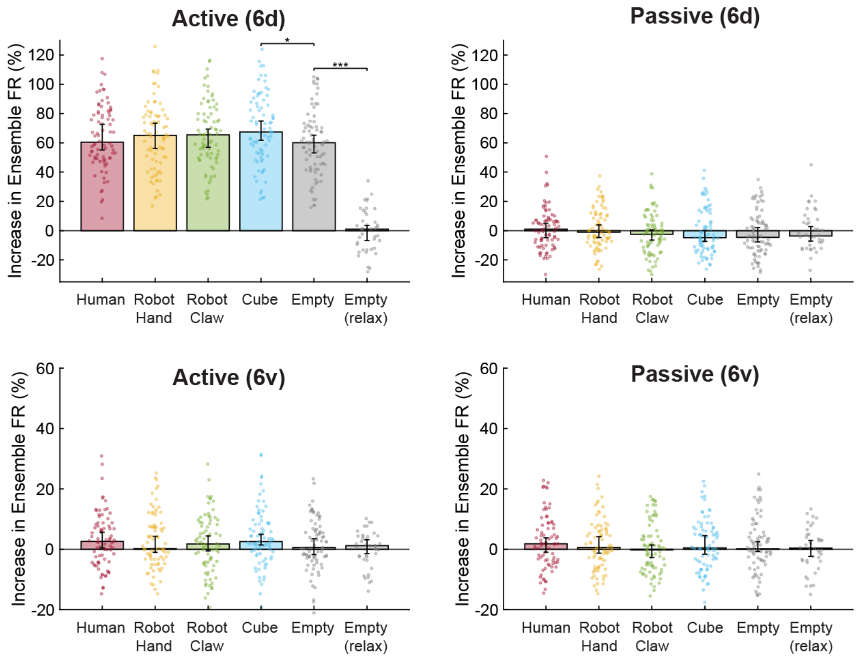


**Figure S9. Modulation of Motor Cortical Ensembles Recorded from T17 6d and 6v Arrays During Virtual Effector Task, Related to Figure 7.**

Increases in ensemble FR during Active (left) and Passive (right) conditions while the participant watched each of the virtual effectors and the baseline condition ("Empty (relax)"). Each point in the swarm plots represents the increase of the mean firing rate of the recorded ensemble (calculated on 1.5s window starting 200ms after the Go Cue) as a percentage of the block baseline FR for one trial. Block baselines were calculated as the mean FR of all the Empty-relax trials (Empty stimulus paired with a "relax" grip cue) in a given block. Bar plots indicate the median increase in ensemble FR, and error bars are 95% confidence intervals (CIs). Outlier trials outside of the 1st-99th percentile range were omitted from each swarm plot, but included in statistical analyses. One-tailed Wilcoxon Rank Sum tests were performed between adjacent conditions to assess whether neural modulation decreased between effector conditions. Asterisks indicate significant $p$-values (*, $p < 0.05$; **, $p < 0.01$; ***, $p < 0.001$). Nonsignificant comparisons ($p < 0.05$) are not shown.

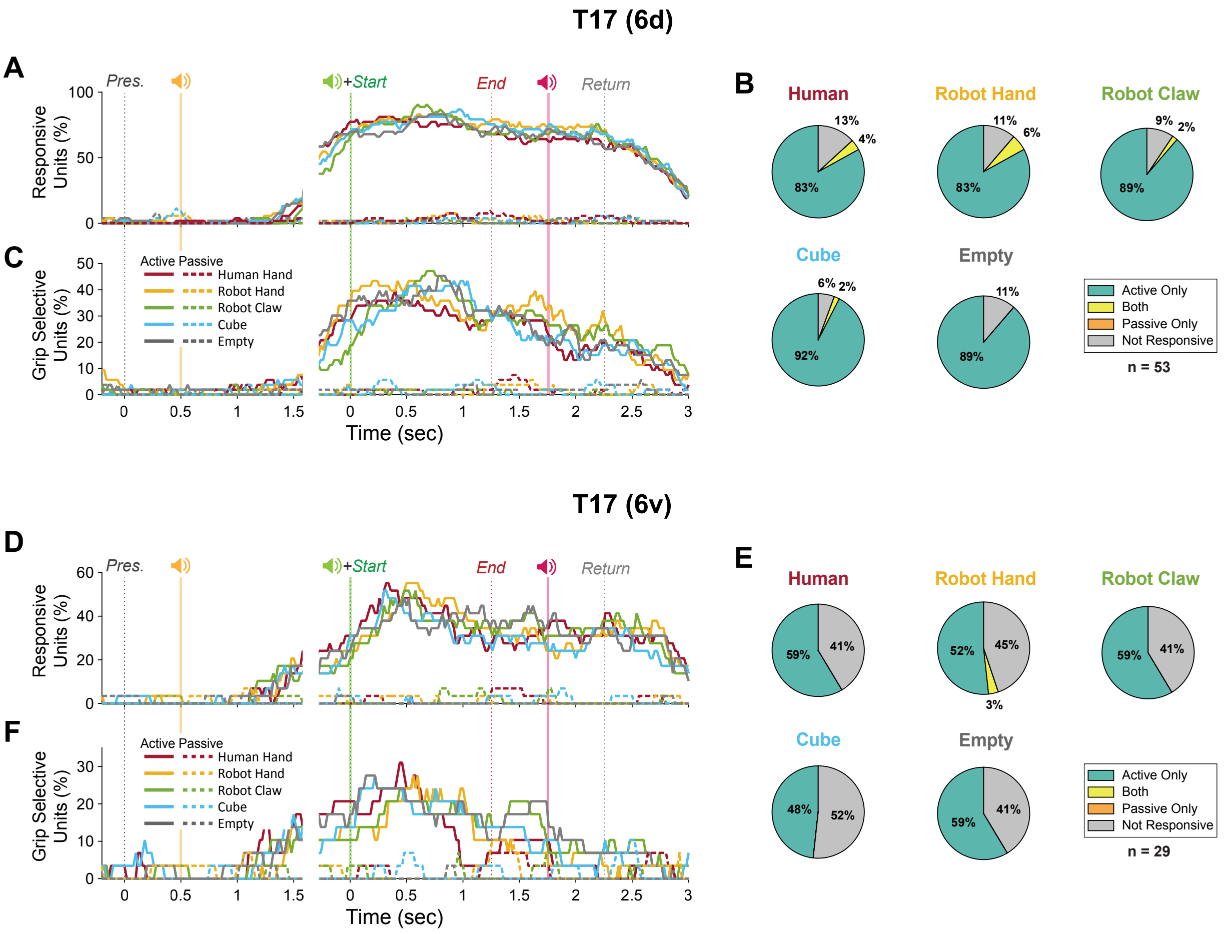


**Figure S10. Responsive and Grip Selective Units During Virtual Effector Task – Participant T17, Related to Figure 7.**

(A) Percent of units from T17's 6d arrays responsive during Active trials (solid lines) and Passive trials (dashed lines) over the course of the trial period for each effector type (see **SI Appendix**).

(B) Percent of units from T17's 6d arrays that were responsive during Active trials, Passive trials, or Both. Data from the time window 200ms-1700ms after the Go Cue was used to evaluate differences in FR between power, pinch, and baseline for each effector and motor intention condition (KW test, $p$ <0.01).

(C) Percent of units from T17's 6d arrays selective for grip type. Units were considered grip selective in a given time window if they had significantly different (RS test, $p < 0.01$) FRs between power and pinch conditions.

(D) Same as (A) but for T17's *6v* arrays.

(E) Same as (B) but for T17's *6v* arrays.

(F) Same as (C) but for T17's *6v* arrays.

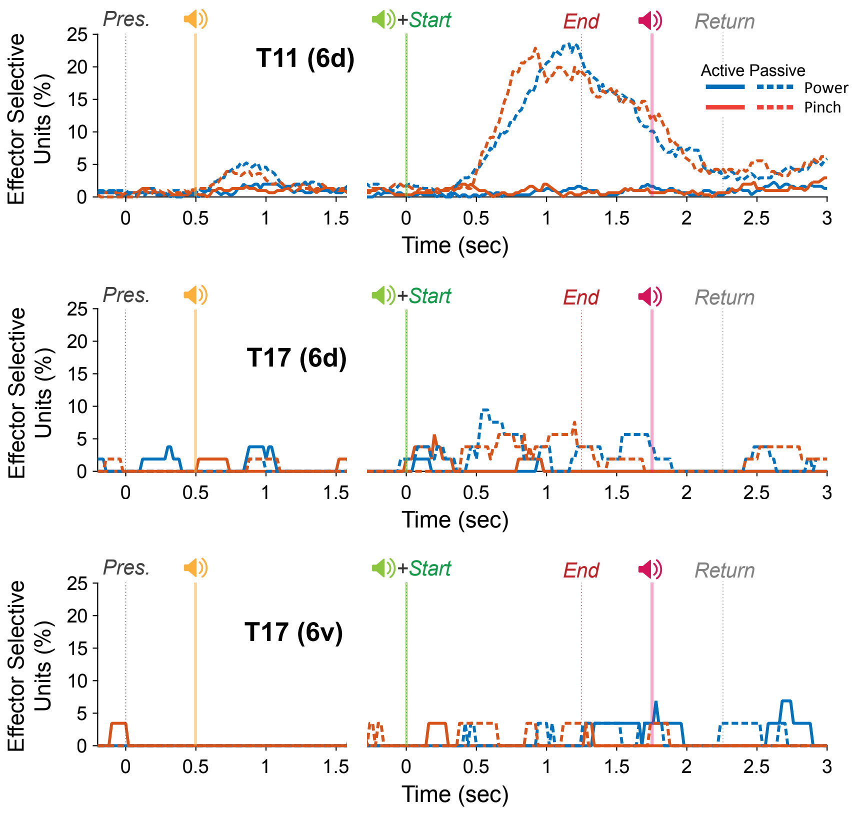


**Figure S11. Percent of Effector Selective Units Over Time in Virtual Effector Task, Related to Figure 7.** Percent of units selective for effector type during Active (solid lines) and Passive (dashed lines) trials. Units were considered effector selective in a given time window if they had significantly different (KW test, $p < 0.01$) firing rates between the five VE Task conditions.

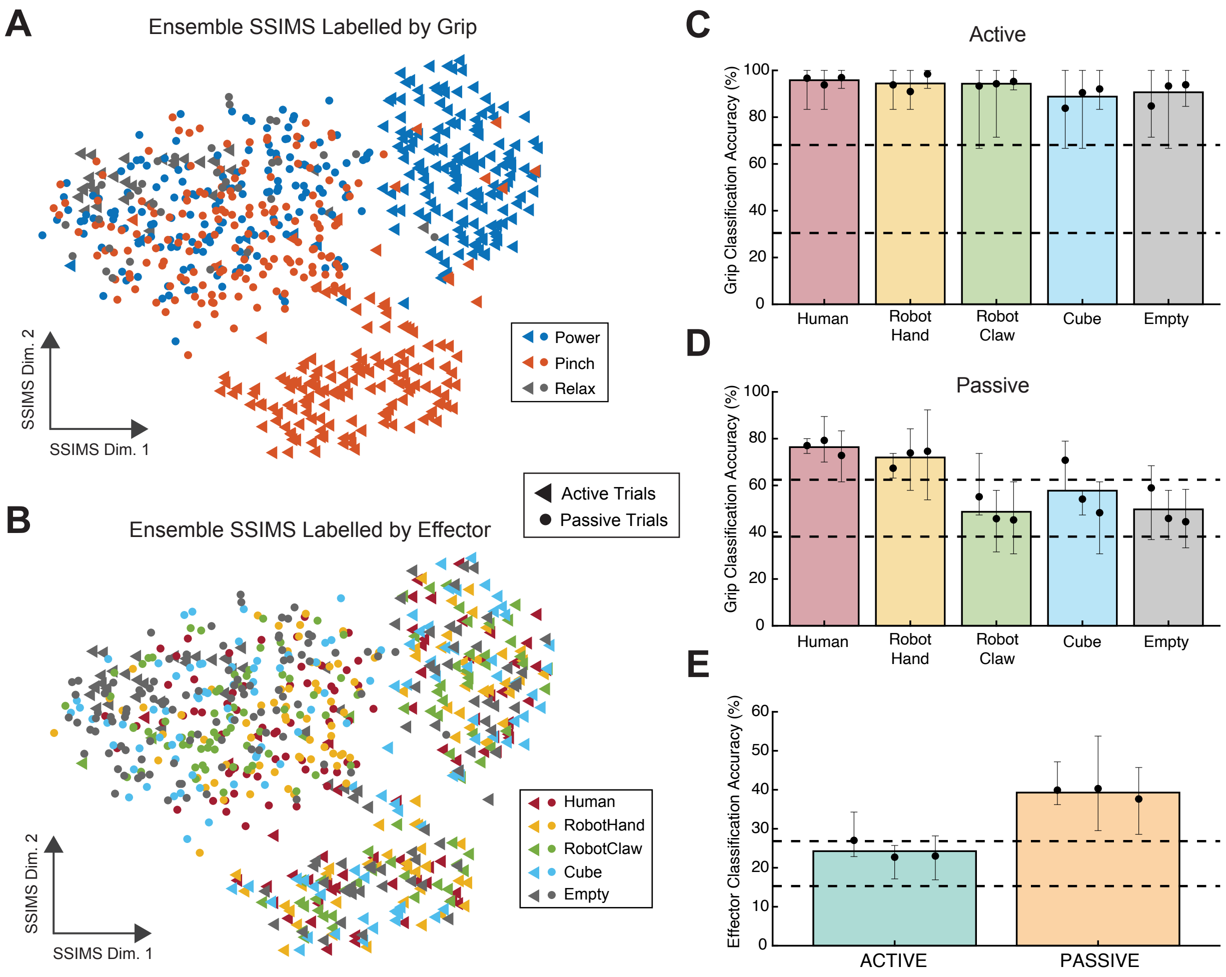


**Figure S12. Ensemble Firing Rate Patterns Reflect Differences between Observed Effector Types during Passive, not Active, Condition, Related to Results Section "Ensemble spiking patterns are similar across effector conditions during Active trials"**

(A-B) 2D spike train similarity space (SSIMS) plots of all trials from a representative VE Task session (T11, trial day 280). Here each marker represents the firing patterns of the entire ensemble of recorded neurons during a single trial. Distance between markers represent the relative similarity between spike train patterns (see **SI Appendix**). Panels (A) and (B) show identical data colored by grip type in (A) and effector type in (B) as indicated in keys.

(C-D) Grip classification accuracies during Active (C) and Passive (D) trials for each effector type. Black dots indicate mean classification performance after 5-fold cross validation for each session. Bar heights denote the mean over all sessions. Error bars denote the range of classification accuracies derived from the cross-validation process for each session. Dashed lines represent the 95% confidence interval of the chance distribution (see **SI Appendix**).

(E) Effector classification accuracies during Active and Passive trials in each session presented similarly as in (C-D).

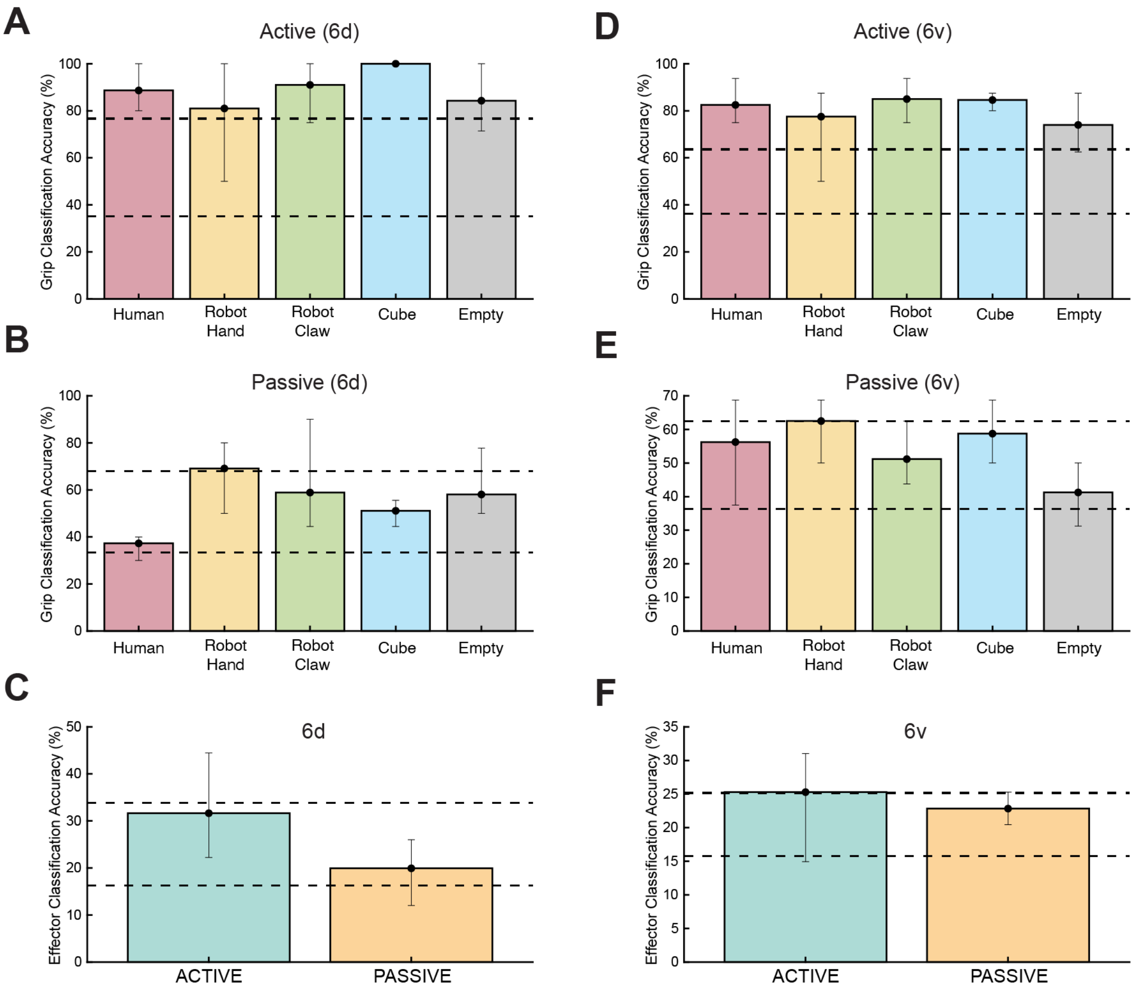


**Figure S13. Grip and Effector Classification of SSIMS Representations of T17 6d and 6v Ensembles During Virtual Effector Task, Related to Results Section "Ensemble spiking patterns are similar across effector conditions during Active trials"**

(A-B) Grip classification accuracies during Active (A) and Passive (B) trials for each effector type applied to 10D SSIMS representations of data from T17's 6d arrays. Black dots and bar heights indicate mean classification performance after 5-fold cross validation. Error bars denote the range of classification accuracies derived from the cross-validation process. Dashed lines represent the 95% confidence interval of the chance distribution (see **SI Appendix**).

(C) Effector classification accuracies on data from T17's 6d arrays during Active and Passive trials presented similarly as in (A).

(D) Same as (A) but for T17's *6v* arrays.

(E) Same as (B) but for T17's *6v* arrays.

(F) Same as (C) but for T17's *6v* arrays.

#### Table S1. Trial and Unit Counts for Each T11 Session, Related to Results Section

| | *VE Task* | | | *Naive JD Task* | *JD Task* | | |
|---|---|---|---|---|---|---|---|
| | Trial Day 262 | Trial Day 276 | Trial Day 280 | Trial Day 372 | Trial Day 358 | Trial Day 387 | Trial Day 412 |
| # of active trials | 176 | 176 | 352 | | | | |
| # of passive trials | 528 | 528 | 352 | 180[a] | 672 | 624 | 768 |
| # of single units | 113 | 90 | 108 | 104 | 110 | 119 | 94 |

[a]only trials from the first 4 blocks of Session 4 are counted (see "**Naive Jittered Dots Task"**)

#### Table S2. Trial and Unit Counts for Each T17 Session, Related to Results Section

| | *JD Task* | *VE Task* |
|---|---|---|
| | Trial Day 62 | Trial Day 90 |
| # of active trials | | 440 |
| # of passive trials | 720 | 440 |
| # of single units (6d) | 86 | 53 |
| # of single units (6v) | 38 | 29 |

**Table S3. Descriptions of the Eight Stimuli Presented in the Jittered Dots Task, Related to Figure 1**

| Effector | Description |
|---|---|
| Human | A human-like hand avatar is used to represent human hand grasps. |
| JD0 | **Dot Starting Positions:** Aligned with the human hand.<br>**Dot Trajectories:** Dots follow the kinematics of the human hand condition. |
| JD1 | **Dot Starting Positions:** Randomly jittered within a range of ±0.25°, vertically and horizontally, relative to the virtual camera.<br>**Dot Trajectories:** same as JD0 |
| JD2 | **Dot Starting Positions:** Randomly jittered within a range of ±1.00°, vertically and horizontally, relative to the virtual camera.<br>**Dot Trajectories:** same as JD0 |
| JD3 | **Dot Starting Positions:** same as JD2<br>**Dot Trajectories:** Same as JD0, but trajectories were randomly rotated within a range of ±45° in all 3 angular dimensions |
| JD4 | **Dot Starting Positions:** same as JD2<br>**Dot Trajectories:** Same as JD0, but trajectories were randomly rotated within a range of ±180° in all 3 angular dimensions (completely random orientation) |
| JD5 | **Dot Starting Positions:** same as JD2<br>**Dot Trajectories:** Replaced with linear translations of the dots in random 3D directions. |
| Empty | A static, white "+" was displayed for the entirety of the trial. |

**Table S4. *P*-values for All Pairwise Comparisons Related to Figure 2.** Wilcoxon Rank Sum test, one-tailed. Significant *p*-values ($p < 0.05$) are bolded. Values above 0.05 indicate no significant difference.

### $6d_{T11}$

*Is ensemble firing rate (FR) when watching this stimulus…*

*…greater than ensemble FR when watching this stimulus?*

| | Human | JD0 | JD1 | JD2 | JD3 | JD4 | JD5 | Empty |
|---|---|---|---|---|---|---|---|---|
| Human | | 0.998 | 1.000 | 1.000 | 1.000 | 1.000 | 1.000 | 1.000 |
| JD0 | **0.002** | | 1.000 | 1.000 | 1.000 | 1.000 | 1.000 | 1.000 |
| JD1 | **6.71e-12** | **8.74e-06** | | 1.000 | 1.000 | 1.000 | 1.000 | 1.000 |
| JD2 | **2.98e-44** | **1.78e-37** | **2.40e-25** | | 0.982 | 0.996 | 1.000 | 1.000 |
| JD3 | **9.84e-50** | **1.62e-43** | **6.16e-33** | **0.018** | | 0.729 | 0.968 | 0.999 |
| JD4 | **9.62e-50** | **5.62e-44** | **5.39e-34** | **0.004** | 0.272 | | 0.888 | 0.997 |
| JD5 | **3.50e-55** | **4.29e-50** | **1.19e-40** | **3.52e-05** | **0.032** | 0.112 | | 0.918 |
| Empty | **3.74e-58** | **5.74e-53** | **1.34e-44** | **5.17e-08** | **6.09e-04** | **0.003** | 0.082 | |

### $6d_{T17}$

*Is ensemble firing rate (FR) when watching this stimulus…*

*…greater than ensemble FR when watching this stimulus?*

| | Human | JD0 | JD1 | JD2 | JD3 | JD4 | JD5 | Empty |
|---|---|---|---|---|---|---|---|---|
| Human | | 1.000 | 1.000 | 1.000 | 1.000 | 1.000 | 1.000 | 1.000 |
| JD0 | **2.22e-05** | | 0.644 | 0.923 | 0.814 | 0.916 | 0.972 | 0.994 |
| JD1 | **4.01e-06** | 0.357 | | 0.912 | 0.721 | 0.901 | 0.949 | 0.994 |
| JD2 | **1.92e-07** | 0.077 | 0.088 | | 0.223 | 0.498 | 0.649 | 0.84 |
| JD3 | **3.26e-07** | 0.186 | 0.280 | 0.778 | | 0.753 | 0.873 | 0.974 |
| JD4 | **2.31e-07** | 0.084 | 0.099 | 0.503 | 0.248 | | 0.666 | 0.823 |
| JD5 | **3.19e-08** | **0.028** | 0.052 | 0.352 | 0.128 | 0.335 | | 0.724 |
| Empty | **5.90e-10** | **0.006** | **0.006** | 0.161 | **0.027** | 0.178 | 0.277 | |

### $6v_{T17}$

*Is ensemble firing rate (FR) when watching this stimulus…*

*…greater than ensemble FR when watching this stimulus?*

| | Human | JD0 | JD1 | JD2 | JD3 | JD4 | JD5 | Empty |
|---|---|---|---|---|---|---|---|---|
| Human | | 0.202 | 0.128 | 0.803 | 0.886 | 0.994 | 0.871 | 1.000 |
| JD0 | 0.799 | | 0.378 | 0.940 | 0.979 | 1.000 | 0.977 | 1.000 |
| JD1 | 0.873 | 0.623 | | 0.976 | 0.990 | 1.000 | 0.989 | 1.000 |
| JD2 | 0.198 | 0.060 | **0.024** | | 0.713 | 0.970 | 0.711 | 1.000 |
| JD3 | 0.115 | 0.021 | **0.010** | 0.288 | | 0.909 | 0.522 | 0.995 |
| JD4 | **0.006** | **3.51e-04** | **7.13e-05** | **0.030** | 0.909 | | 0.090 | 0.896 |
| JD5 | 0.129 | **0.023** | **0.011** | 0.290 | 0.479 | 0.910 | | 0.995 |
| Empty | **9.96e-05** | **1.24e-06** | **9.53e-08** | **3.55e-04** | **0.005** | 0.104 | **0.005** | |

### VIDEO TITLE/LEGENDS

**Video S1. Jittered Dots Task Stimuli, Related to Figure 1.**
Videos of the eight stimulus types used during the Jittered Dots (JD) task. The participant was instructed to provide a numeric rating (1 to 4) of the perceived anthropomorphicity (ARS rating) of each pseudorandomly presented stimulus (**Figure 1B**).

**Video S2. The "Aha" Moment, Related to Figure 6.**
Mean ensemble firing rate (FR) during the Naive Jittered Dots Task before and after T11 said he "figured out" that the dot stimuli depicted hand-like grasping movements. Unit FRs were calculated in 20 ms bins, smoothed with an 80 ms Gaussian kernel, and averaged (mean) across all units. Plotted traces represent this ensemble FR smoothed using a 1 second moving average window. Real-time ensemble FR traces are plotted over the trial-averaged responses of all trials before the "Aha!" Moment (left) and all trials after the "Aha!" Moment (right). Shaded error bars represent 95% confidence intervals.

**Video S3. Virtual Effector Task Stimuli, Related to Figure 7.**
Videos of the five "effectors" shown during the Virtual Effector (VE) Task.

#### Supplementary Discussion 1: Anthropomorphic Tuning as an Indicator of Effector Predictability

In addition to differences of the neural responses during the movement phase of the JD Task, we also observed that the most anthropomorphic conditions evoked neuronal responses even before the start of movement (**Figure 2C**). Much like previous fMRI studies that showed neural responses in the motor cortex to the presentation of static images of human movement[8–10], a subset of recorded neurons appeared to be responsive to the simple appearance of the Human, JD0, and (to a lesser extent) JD1 stimuli at the beginning of the trial (**Figure 2C**). Furthermore, there was anticipatory ramping of neuronal activation prior to the start of stimulus movement during the Human and JD0 conditions that is similar to what has been found among F5 neurons in monkeys during action observation[11]. Notably, observation-related anticipatory activity in F5 neurons were found only when monkeys observed hand grasps in a predictable context[11]; when monkeys observed an experimenter perform hand grasps at random, there was no anticipatory activation among F5 neurons[12].

Evidence of neuronal activation of F5 neurons prior to the start of observed movement has been interpreted[13] as support for predictive coding accounts of the "mirror neuron system"[14–16], which model mirror activity in the cortex as a predictive system as opposed to a strictly feedforward pattern recognizer[17]. The predictive coding model of mirror activity posits that action understanding could theoretically be carried out through the comparison of forward models of others' actions with incoming sensory information at several levels of abstraction distributed across the brain. In this framework, the premotor cortex, which is already specialized for representing motor plans during - and prior to - execution, might be responsible for comparing observed movement patterns to models of expected movements based on context cues arising from the prefrontal cortex. The involvement of PMd in generating predictions of others' movements has since been shown in human fMRI[18] and monkey[19] studies, and recent studies have demonstrated that impairment of PMd due to rTMS[20] or prior injury[21] can significantly diminish performance in motor prediction tasks. Could our results reflect activity related to motor prediction in the human hand knob area?

In their MOSAIC model for motor learning and control, Haruno et al.[22] proposed that motor control can be modeled as a collection of controllers, each with a pair of forward and inverse models of movement that are selected according to a "responsibility signal" representing the relative involvement that controller should have in the given context. Wolpert et al.[23] extended their model to help explain action observation; the brain simultaneously runs multiple forward models predicting the behavior of the individual they are observing and compares these with sensory inputs to generate *responsibilities* representing the degree to which their internal models explain the observed behavior. Likewise, it is possible that anthropomorphicity-related changes in activation that we observed in the surface of the human hand knob area could correspond to the relative involvement of the innate motor system in predicting movement in the observed stimuli. In other words, contextual cues like prior knowledge (**Figure 6**) and effector appearance before movement could combine with sensory feedback during effector movement to generate error signals that modulate the weighting, or *responsibility*, of the internal motor model in the task of predicting sensory consequences in the observed movement (and thus, perhaps, in accomplishing the instructed task of determining whether the stimulus was hand-like or not). This interpretation could help explain the large difference in neuronal activation we observed between JD1 and JD2 in $6d_{T11}$ (**Figure 2C**). Whereas the starting appearances of the Human, JD0, and JD1 stimuli were predictably tied to particular movement patterns, the starting appearance of JD2 was identical to that of JD3, JD4, and JD5. Thus, the relative utility of the internal motor model in generating sensory predictions could not be assessed until after movement started. We observed a separation between neuronal activation traces JD2-5 (**Figure 2C**) only after movement began (~500ms), which might indicate the point at which enough information was accumulated to determine whether the dots moved in a predictable, human-like manner, and warranted additional motor cortical recruitment (e.g. JD2) or not (e.g. JD5). Meanwhile, early contextual cues offered by the most anthropomorphic conditions (Human, JD0, JD1) could have biased the responsibility signal toward the internal model of hand movement (instead of, say, the internal model of random dot movement) allowing for a priming of the network that resulted in a more extensive neural response once movement actually began. The contribution of network "priming" could also help explain why neural responses to JD0 and JD1 in participant T17 were less robust than in T11 (**Figure 2**). Participant T11, who performed three times as many trials of the JD Task as T17 (**Tables S1-2**), had more of an opportunity to learn the association between the static, starting images of JD0 and JD1 with the resultant anthropomorphic hand movement.

Of course, this explanation for the observed anthropomorphicity gradient is one of several that could be inferred from our current results. Future iBCI studies that specifically employ movement prediction tasks would better elucidate what role, if indeed any, the motor cortex might play in action prediction or understanding.

#### Supplementary Discussion 2: Grip Separability During Active Condition

Although grip classification accuracies were not significantly different across VE Task effectors during the Active condition (**Figure S12C**), it is notable that average accuracy values were slightly lower for the least anthropomorphic effectors (Cube: 89%, Empty: 91%) compared to the more anthropomorphic effectors (Human: 96%, Robot Hand: 95%, Robot Claw: 95%). Additionally, traces of the percentage of grip selective units in $6d_{T11}$ over time appear to show less grip selectivity during Cube and Empty trials compared to other effectors during attempted movement (**Figure 7E**, bottom). These results suggest that the neural responses from attempted power and pinch trials were slightly more distinct, or separable, when the participant was viewing more anthropomorphic effectors. However, this result is difficult to interpret, because although T11 was instructed to use the audio cues (e.g. "power") to determine what grip to attempt, it is unclear what effect the anthropomorphic congruence of the visual stimulus had on his ability to consistently perform the instructed motor imagery. For example, the visual feedback of the more anthropomorphic effectors may have inadvertently served as a second "grip cue" that the participant utilized to ensure he was performing the correct motor imagery, resulting in "cleaner" grip attempts and less variability in the neural data for those trials. This highlights the complexity of assessing the effects of task-relevant sensory input on iBCI control, as it is often difficult to isolate the neural effects of sensory input to the brain from the semantic contribution that input could have on performance of the task.

### SUPPLEMENTAL REFERENCES

1. Glasser, M. F. *et al.* A multi-modal parcellation of human cerebral cortex. *Nature* **536**, 171–178 (2016).
2. Glasser, M. F. *et al.* The minimal preprocessing pipelines for the Human Connectome Project. *Neuroimage* **80**, 105–124 (2013).
3. Simeral, J. D. *et al.* Home use of a percutaneous wireless intracortical brain-computer interface by individuals with tetraplegia. *IEEE Trans. Biomed. Eng.* **68**, 2313–2325 (2021).
4. Vargas-Irwin, C. & Donoghue, J. P. Automated spike sorting using density grid contour clustering and subtractive waveform decomposition. *J. Neurosci. Methods* **164**, 1–18 (2007).
5. Vargas-Irwin, C. E., Brandman, D. M., Zimmermann, J. B., Donoghue, J. P. & Black, M. J. Spike train SIMilarity Space (SSIMS): a framework for single neuron and ensemble data analysis. *Neural Comput.* **27**, 1–31 (2015).
6. Victor, J. D. & Purpura, K. P. Nature and precision of temporal coding in visual cortex: a metric-space analysis. *J. Neurophysiol.* **76**, 1310–1326 (1996).
7. van der Maaten, L. & Hinton, G. Visualizing data using t-SNE. *J. Mach. Learn. Res.* **9**, 2579–2605 (2008).
8. Hamzei, F. *et al.* The human action recognition system and its relationship to Broca's area: an fMRI study. *Neuroimage* **19**, 637–644 (2003).
9. Quandt, L. C., Lee, Y.-S. & Chatterjee, A. Neural bases of action abstraction. *Biol. Psychol.* **129**, 314–323 (2017).
10. Assmus, A., Giessing, C., Weiss, P. H. & Fink, G. R. Functional interactions during the retrieval of conceptual action knowledge: an fMRI study. *J. Cogn. Neurosci.* **19**, 1004–1012 (2007).
11. Maranesi, M., Livi, A., Fogassi, L., Rizzolatti, G. & Bonini, L. Mirror neuron activation prior to action observation in a predictable context. *J. Neurosci.* **34**, 14827–14832 (2014).
12. Maranesi, M. *et al.* Monkey gaze behaviour during action observation and its relationship to mirror neuron activity. *Eur. J. Neurosci.* **38**, 3721–3730 (2013).
13. Urgen, B. A. & Miller, L. E. Towards an empirically grounded predictive coding account of action understanding. *J. Neurosci.* **35**, 4789–4791 (2015).
14. Kilner, J. M., Friston, K. J. & Frith, C. D. Predictive coding: an account of the mirror neuron system. *Cogn. Process.* **8**, 159–166 (2007).
15. Kilner, J. M. More than one pathway to action understanding. *Trends Cogn. Sci.* **15**, 352–357 (2011).
16. Friston, K., Mattout, J. & Kilner, J. Action understanding and active inference. *Biol. Cybern.* **104**, 137–160 (2011).
17. Giese, M. A. & Poggio, T. Neural mechanisms for the recognition of biological movements. *Nat. Rev. Neurosci.* **4**, 179–192 (2003).
18. Stadler, W. *et al.* Predicting and memorizing observed action: differential premotor cortex involvement. *Hum. Brain Mapp.* **32**, 677–687 (2011).
19. Cirillo, R., Ferrucci, L., Marcos, E., Ferraina, S. & Genovesio, A. Coding of self and other's future choices in dorsal premotor cortex during social interaction. *Cell Rep.* **24**, 1679–1686 (2018).
20. Brich, L. F. M., Bächle, C., Hermsdörfer, J. & Stadler, W. Real-time prediction of observed action requires integrity of the dorsal premotor cortex: evidence from repetitive transcranial magnetic stimulation. *Front. Hum. Neurosci.* **12**, 101 (2018).
21. de Wit, M. M. & Buxbaum, L. J. Critical motor involvement in prediction of human and non-biological motion trajectories. *J. Int. Neuropsychol. Soc.* **23**, 171–184 (2017).
22. Haruno, M., Wolpert, D. M. & Kawato, M. Mosaic model for sensorimotor learning and control. *Neural Comput.* **13**, 2201–2220 (2001).
23. Wolpert, D. M., Doya, K. & Kawato, M. A unifying computational framework for motor control and social interaction. *Philos. Trans. R. Soc. Lond. B Biol. Sci.* **358**, 593–602 (2003).
